# Experimental evaluation of thermodynamic speed limit in living cells via information geometry

**DOI:** 10.1101/2020.11.29.403097

**Authors:** Keita Ashida, Kazuhiro Aoki, Sosuke Ito

**Affiliations:** Universal Biology Institute, The University of Tokyo, 7-3-1 Hongo, Bunkyo-ku, Tokyo 113-0033, Japan; Quantitative Biology Research Group, Exploratory Research Center on Life and Living Systems (ExCELLS), National Institutes of Natural Sciences, 5-1 Higashiyama, Myodaiji-cho, Okazaki, Aichi 444-8787, Japan; Division of Quantitative Biology, National Institute for Basic Biology, National Institutes of Natural Sciences, 5-1 Higashiyama, Myodaiji-cho, Okazaki, Aichi 444-8787, Japan; Department of Basic Biology, School of Life Science, SOKENDAI (The Graduate University for Advanced Studies), 5-1 Higashiyama, Myodaiji-cho, Okazaki, Aichi 444-8787, Japan; Department of Physics, The University of Tokyo, 7-3-1 Hongo, Bunkyo-ku, Tokyo 113-0033, Japan; JST, PRESTO, 4-1-8 Honcho, Kawaguchi, Saitama, 332-0012, Japan

## Abstract

Chemical reactions are responsible for information processing in living cells, and thermodynamic trade-off relations can explain their accuracy and speed. Its experimental test in living cells had not existed despite its importance because it is hard to justify sample size sufficiency. This paper reports the first experimental test of the thermodynamic trade-off relation, namely the thermodynamic speed limit, in living systems at the single-cell level where the sample size is relatively small. Due to the information-geometric approach, we can demonstrate the thermodynamic speed limit for the extracellular signal-regulated kinase phosphorylation using time-series fluorescence imaging data. Our approach quantifies the intrinsic speed of cell proliferation and can potentially apply other signal transduction pathways to detect their information processing speed.

**One-Sentence Summary:** Experimental measurement of information thermodynamic speed by fluorescence imaging in living cells

## Main Text

The Gibbs free energy change mainly drives information transmission in living cells, and its mechanism follows thermodynamic laws. For example, systems biology reveals a deep connection between information transmission accuracy and the Gibbs free energy change in a cell *(1)*. In recent developments of stochastic thermodynamics *(2)* and chemical thermodynamics *(3, 4)*, a similar connection occurs more deeply. The trade-off relations among speed, thermodynamic cost, and accuracy in living cells have been proposed *(5, 6)*, and its general thermodynamic bound of information processing in signal transduction has been discussed in terms of Maxwell’s demon *(7–10)* and thermodynamic generalizations of uncertainty relations *(11–14)*. Especially, as a generalization of the quantum speed limit, thermodynamic speed limits have been proposed in several ways *(13–22)*, and some variants of thermodynamic speed limits are based on the mathematical properties of the Fisher information *(13, 15, 17–19, 21, 22)*, which gives a metric of information geometry *(23)*. Historically, information geometry has been considered as a possible choice of differential geometry for equilibrium thermodynamics *(24, 25)*. In recent years, information geometry meets nonequilibrium thermodynamics such as stochastic thermodynamics *(13, 19, 26)* and chemical thermodynamics *(22)*, and it provides a unified framework to derive these trade-off relations in living systems. Despite thermodynamic trade-off relations has been proposed for biological applications, an experimental test of thermodynamic trade-off relations in living systems has not yet been reported. Moreover, the thermodynamic speed limits have not been examined even in artificial systems, while the thermodynamic uncertainty relations have been examined in artificial systems *(27–29)*.

This paper experimentally examined the thermodynamic speed limit by quantifying the Fisher information of time from the fluorescence imaging of the extracellular signal-regulated kinase (ERK) phosphorylation in normal rat kidney epithelial (NRK-52E) cells. In information geometry, the Fisher information of time is regarded as the square of the intrinsic speed. This Fisher information is calculated from the time evolution of the phosphorylated ERK fraction, which can be experimentally measured by the Förster resonance energy transfer (FRET) signal. We also evaluated other information-geometric quantities for activation and inactivation processes from this Fisher information. These information-geometric quantities illustrate the thermodynamic cost of the ERK phosphorylation, which is related to the Gibbs free energy change and the change speed of the observable. We focused on the thermodynamic speed limit based on these information-geometric quantities and found that the speed limit’s efficiency ranges from 0.4 to 0.9 for the ERK phosphorylation in living NRK-52E cells. We quantified the thermodynamic variability for cell density changes and the perturbation of the upstream Raf pathway. While the cell density increases the thermodynamic cost and reduces the efficiency, the Raf inhibitor addition reduces the thermodynamic cost, but the efficiency is robust to the Raf inhibitor addition.

The information-geometric method proposed in this paper is generally applicable to switching dynamics between the active state and the inactive state. We consider the following chemical reaction

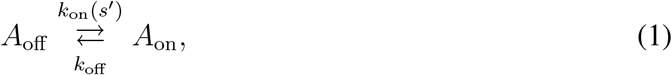

where *A*_on_ is an active state, *A*_off_ is an inactive state, *s*′ is stimulus, and *k*_on_*(s*′) and *k*_off_ are rate constants. Because *k*_on_*(s*′) can depend on stimulus *s*′, it causes a pulsatile response of *A*_on_ if this stimulus *s*′ is excitable. This paper focuses on the ERK phosphorylation in the Ras-Raf-MEK-ERK pathway that shows a pulsatile response *(30, 31)*. In Fig. 1A, we show the Ras-Raf-MEK-ERK pathway that relays extracellular stimuli such as growth factors from the plasma membrane to targets in the cytoplasm and nucleus. This three-tiered Raf-MEK-ERK mitogen-activated protein kinase (MAPK) cascade plays an essential role in various cellular processes, including cell proliferation, differentiation, and tumorigenesis *(32)*. Upon growth factor stimulation, the receptor tyrosine kinase (RTK) activates the Ras small GTPase at the plasma membrane, which recruits and activates the Raf. The activated Raf induces activation and phosphorylation of the MEK. The upstream kinase MEK phosphorylates the ERK to increase kinase activation of the ERK. The phosphorylated ERK is finally dephosphorylated by phosphatases, thereby shutting down the ERK activation. The phosphorylated MEK catalyzes a phosphate transfer from the adenosine triphosphate (ATP) to the ERK, and the Gibbs free energy difference of ATP hydrolysis thermodynamically drives this phosphorylation of the ERK *(32)*. The phosphorylated ERK directly phosphorylates proteins such as the ELK1 and ribosomal S6 kinase (RSK), which induce the expressions of transcriptional factors including EGR and c-Fos *(31–33)*. These transcriptional factors promote gene expression for cell cycle progression, and thus the ERK phosphorylation regulates cell proliferation. Here, *A*_on_ corresponds to the phosphorylated state of the ERK, *A*_off_ corresponds to the nonphosphorylated state of the ERK, and *s*′ corresponds to the stimulus by upstream proteins in the Ras-Raf-MEK-ERK pathway, respectively. We try to examine the thermodynamic speed limit for the ERK phosphorylation in living cells using the time series of the phosphorylated ERK fraction, which can be experimentally measured by the FRET technique at the single-cell level *(31, 34)*.

**Fig. 1.**
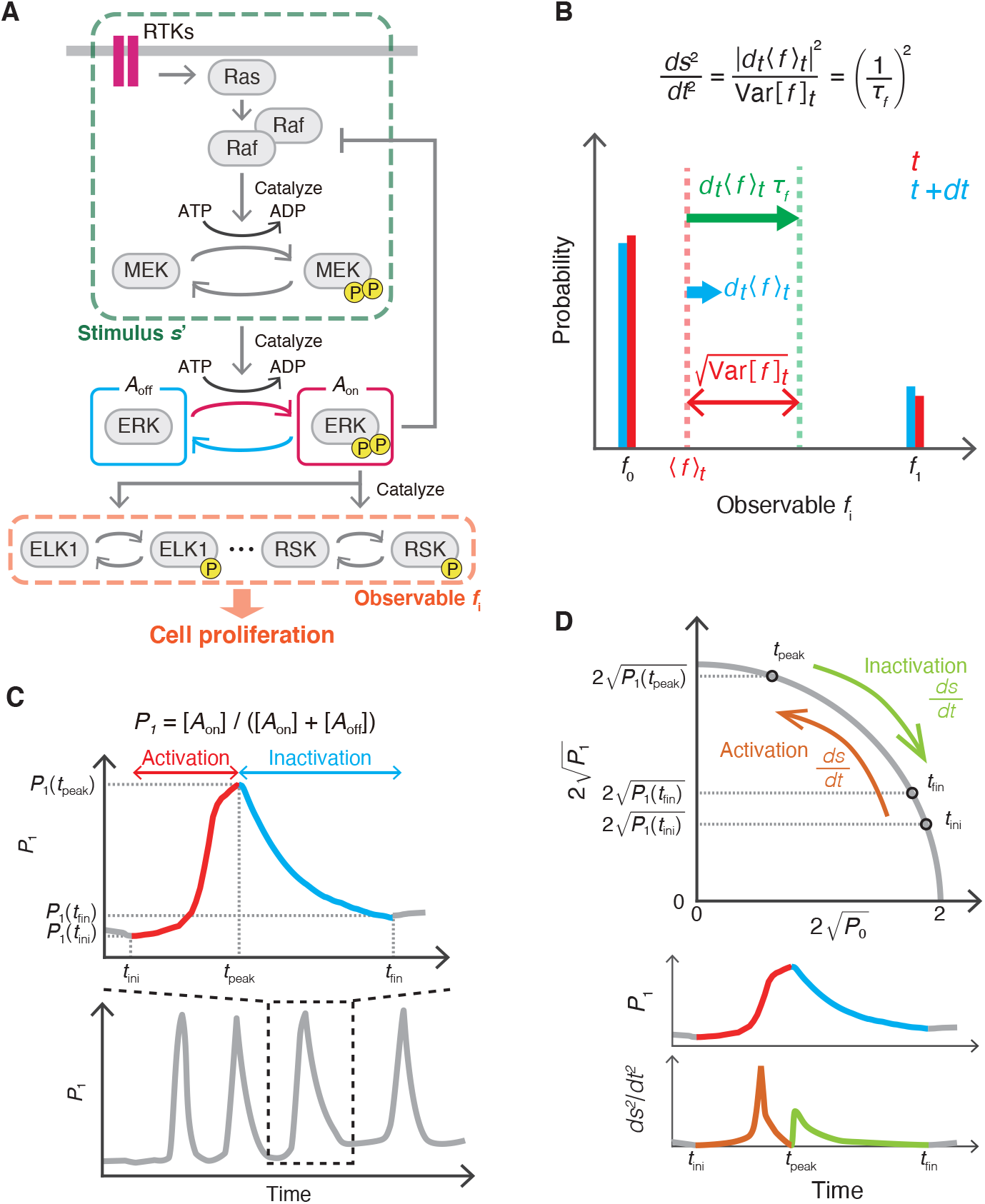
Schematic of the ERK phosphorylation and information geometry. (**A**) The Ras-Raf-MEK-ERK pathway as an example of switching dynamics between the active state *A*_on_ and the inactive state *A*_off_ . The upstream proteins correspond to the stimulus *s*′, and the Raf on the upstream pathway affects the phosphorylated ERK. The phosphorylated ERK directly catalyzes the ELK1 and RSK phosphorylation, which are possible candidates of the observable *f*_*i*_. (**B**) The intrinsic speed *ds/dt* corresponds to 1*/τ*_*f*_, which implies the change speed of the observable *f*_*i*_. The expected value of observable *f*_*i*_ varies from its current value ⟨ *f* ⟩ _*t*_ to 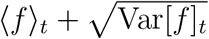 during time interval *τ*_*f*_ if the speed |*d*_*t*_ ⟨ *f* ⟩_*t*_ | is constant. (**C**) Typical behavior of the phosphorylated ERK fraction *P*_1_ in the activation and inactivation processes. The time *t*_ini_ *(t*_peak_) indicates the beginning of the activation (inactivation), and the time *t*_peak_ *(t*_fin_) indicates the end of the activation (inactivation). (**D**) Schematic of the manifold in information geometry and the intrinsic speed *ds/dt* on this manifold. The activation and inactivation processes give at least two peaks in the time series of *ds*^2^*/dt*^2^.

We discuss the thermodynamic cost estimation from the time series of the phosphorylated ERK fraction *P*_1_ = [*A*_on_]*/*([*A*_on_] + [*A*_off_]), where [*A*_on_] and [*A*_off_] are concentrations corresponding to *A*_on_ and *A*_off_, respectively (see also supplementary materials). Because the total concentration [*A*_tot_] = [*A*_off_] + [*A*_on_] is conserved, the nonphosphorylated and phosphorylated ERK fractions *P*_0_ = 1 − *P*_1_ = [*A*_off_]*/*[*A*_tot_] and *P*_1_ = [*A*_on_]*/*[*A*_tot_] can be regarded as the probability distribution, and a Riemannian manifold can be introduced as the set of probability distributions in information geometry *(23)*. In general, it is hard to estimate the Gibbs free energy change in living cells because it needs prior knowledge about the equilibrium concentration *(3)* and too large sample sizes to estimate the rate constants experimentally. However, this equilibrium concentration is not estimated well from the oscillating time series of *P*_1_, and the sample size of live-cell imaging data should be small. We here propose a novel information-geometric method to estimate the thermodynamic cost from the time series of *P*_1_. This method provides the estimation of the thermodynamic cost from the single shot of the live-cell imaging data and does not require prior knowledge about the equilibrium concentration. This method focuses on an intrinsic speed 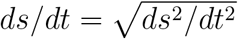 on the manifold of information geometry, which is related to the thermodynamic cost such as the Gibbs free energy change and the speed of the observable associated with the ERK phosphorylation *(13, 19, 22)*. The square of this intrinsic speed is defined as the Fisher information of time

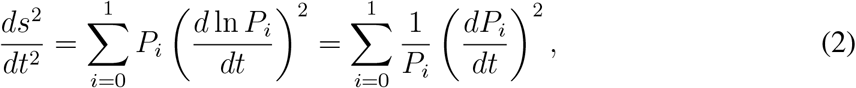

which can be estimated from the time series of *P*_1_ because *ds*^2^*/dt*^2^ only consists of the concentration fraction *P*_*i*_ and its change speed *dP*_*i*_*/dt*.

The square of the intrinsic speed gives a fundamental speed of dynamics of the observable related to the ERK phosphorylation. Let *f*_*i*_ be an observable of the ERK states, where *i* = 0 *(i* = 1) means the nonphosphorylated (phosphorylated) state (Fig. 1A and 1B). Because any observable of a binary state *f*_*i*_ can be regarded as an efficient estimator, the square of the intrinsic speed *ds*^2^*/dt*^2^ is given by a consequence of the Cramér-Rao bound for an efficient estimator *(23)*

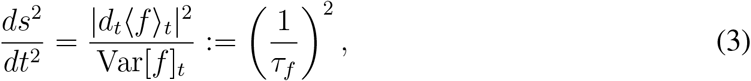

where ⟨*f* ⟩_*t*_ = ∑_*i*_ *P*_*i*_*(t)f*_*i*_ is the mean value, |*d*_*t*_⟨*f* ⟩_*t*_| = |*d*⟨*f* ⟩_*t*_*/dt*| implies the expected change speed of the observable, and 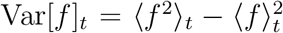 indicates the observable fluctuation. Because the expected value of the observable varies from its current value ⟨*f* ⟩_*t*_ to the standard deviation 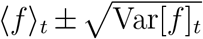 during the time interval *τ*_*f*_, the intrinsic speed *ds/dt* = 1*/τ*_*f*_ implies the change speed of any observable *f*_*i*_. Possible candidates of the observable *f*_*i*_ in the Ras-Raf-MEK-ERK pathway are the phosphorylation fractions of the ELK1 and RSK because the phosphorylated ERK directly catalyzes their phosphorylation (Fig. 1A). These phosphorylation fractions regulate cell proliferation, and thus the intrinsic speed provides the regulation speed of cell proliferation. We stress that there is a deep connection between the intrinsic speed and the entropy production rate (see also supplementary materials). In thermodynamics, the entropy production rate *σ* = − *(dG/dt)/T* is the fundamental cost of dynamics, where *G* is the Gibbs free energy and *T* is the temperature of the medium. Because the entropy production rate is related to the speed of the Gibbs free energy change, a relation between the entropy production rate and the intrinsic speed holds for a binary state in parallel with Eq. (3),

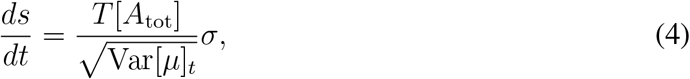

where *µ*_0_ and *µ*_1_ are the chemical potentials of *A*_off_ and *A*_on_, respectively. Therefore, the intrinsic speed is proportional to the entropy production rate of ERK phosphorylation *σ*. Moreover, the Fisher information of time is also related to the time derivative of the entropy production rate under near-equilibrium conditions *ds*^2^*/dt*^2^ ≃ − *(dσ/dt)/*(2*R*[*A*_tot_]), where *R* is the gas constant *(22)*.

As shown in Fig. 1C, a pulsatile response consists of activation and inactivation processes, and we introduce information-geometric quantities of these processes as measures of the thermodynamic cost and the speed limit’s efficiency (see also supplementary materials). During the activation (inactivation) process, the phosphorylated ERK fraction *P*_1_ is monotonically increasing (decreasing) in time. The activation (inactivation) starts at time *t*_0_ = *t*_ini_ *(t*_0_ = *t*_peak_) and ends at time *t*_1_ = *t*_peak_ *(t*_1_ = *t*_fin_). We here introduce three information-geometric quantities *(13, 25)*, the action

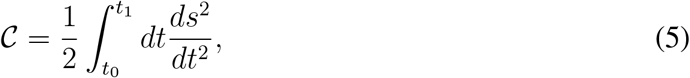

the length

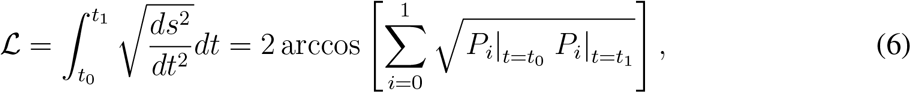

and the mean velocity

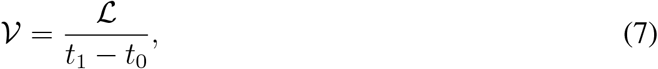

during the process from time *t*_0_ to *t*_1_. Because the intrinsic speed *ds/dt* is related to the ratio between the entropy production rate *σ* and the standard deviation of the chemical potential 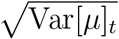 in Eq. (4), these geometric quantities 𝒞, ℒ, 𝒱 reflect thermodynamic properties of activation and inactivation processes. Biologically, these quantities also reflect the regulation of cell proliferation in terms of Eq. (3). Moreover, the action 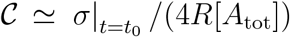 is approximately proportional to the entropy production rate at time *t*_0_ under near-equilibrium conditions *(22)*. Thus, the 𝒞 action can be regarded as the thermodynamic cost of this process. The length ℒ is given by twice the Bhattacharyya angle, which is a measure of a difference between two concentration fractions at time *t*_0_ and time *t*_1_. This length ℒ is also known as the thermodynamic length *(25)*, which is a measure of thermodynamics for finite time transformations. The mean velocity 𝒱 quantifies the speed of the concentration fraction change during the process. Because the mean velocity is the time-averaged intrinsic speed, it quantifies the time-averaged regulation speed of cell proliferation.

In information geometry, ℒ is regarded as the arc length of a circle in 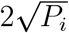 coordinate, and *ds/dt* is the intrinsic speed on this circle (see also Fig. 1D). From the Cauchy-Schwarz inequality, we obtain the thermodynamic speed limit *(13)*

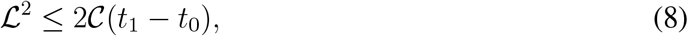

which is a trade-off relation between the thermodynamic cost 𝒞 and the transition time *t*_1_ −*t*_0_ during the process. To quantify how much the thermodynamic cost converts into the concentration fraction change speed, we can consider the speed limit’s efficiency *(13)*

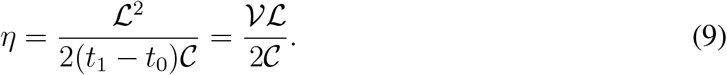

The efficiency *η* satisfies 0 ≤ *η* ≤ 1, and *η* = 1 *(η* = 0) implies that this conversion is most efficient (inefficient). The efficiency becomes higher if the intrinsic speed is close to constant because *η* = 1 if and only if the intrinsic speed *ds/dt* is constant regardless of time, *(d/dt)* |*ds*^2^*/dt*^2^ |= 0.

We experimentally measured the phosphorylated ERK fraction in living NRK-52E cells with the FRET-based ERK biosensor, the EKAREV-NLS (Fig. 2A) *(34)*. By comparing the fluorescence ratio of cells with the phosphorylated ERK fraction obtained from the western blotting (Fig. 2B), we quantified *P*_1_ and the square of the intrinsic speed *ds*^2^*/dt*^2^ for the ERK activation *(p* = a) and inactivation *(p* = i) processes under the condition of different cell densities: 2.0 × 10^3^ cells/cm^2^ (low, *d* = L), 2.0 × 10^4^ cells/cm^2^ (medium, *d* = M), and 2.0 × 10^5^ cells/cm^2^ (high, *d* = H), where the indices *p* ∈ {a, i} and *d* ∈ {L, M, H}regard the process and cell density, respectively. The pulses of the ERK activation were observed under these conditions (Fig. 2C and movie S1), and the behavior of *ds*^2^*/dt*^2^ characterizes these conditions *p* and *d* (Fig. 2D). It reveals thermodynamic differences of the intracellular ERK activation under these conditions.

**Fig. 2.**
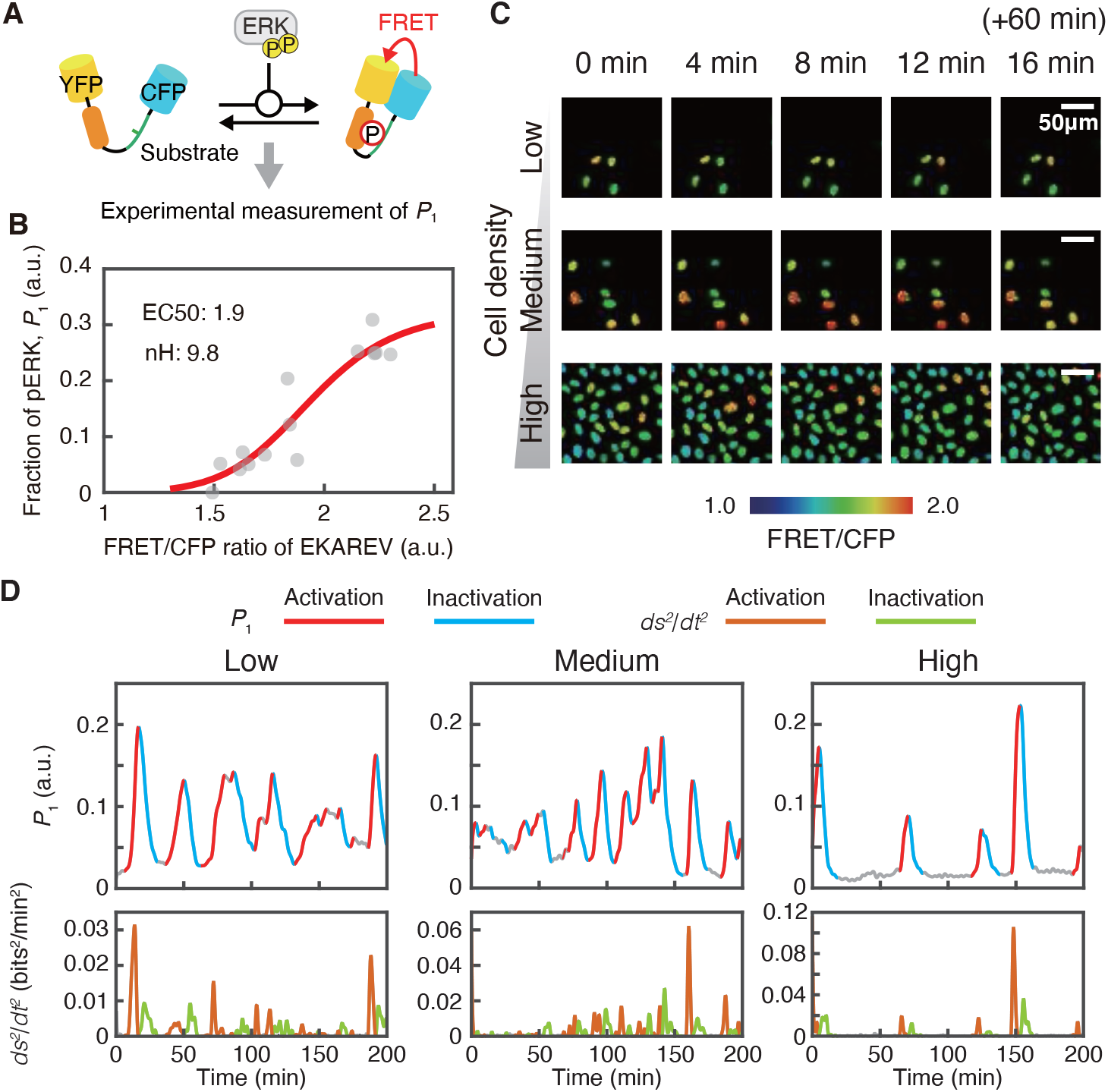
The dynamics of the ERK phosphorylation under the conditions of different cell densities. (**A**) Schematic of the FRET-based ERK sensor. The phosphorylated ERK fraction *P*_1_ can be measured from this FRET signal. (**B**) The relation between the FRET/CFP ratios of the EKAREV-NLS and the phosphorylated ERK fraction *P*_1_. The red line indicates the fitted curve of the Hill equation. (**C**) Representative images of the phosphorylated ERK under the conditions of low, medium, and high cell densities. The time 0 min implies the timing that one hour has passed since imaging initiation. (**D**) Representative examples of the ERK phosphorylation dynamics under low (left), medium (middle), and high (right) cell densities. The unit (a.u.) indicates the arbitrary unite. This graph shows the time series of *P*_1_ and *ds*^2^*/dt*^2^ from imaging initiation. We use bits (which is the unit called shannon, Sh) as the unit of information where 1 (bits) = ln 2 (nats). The Fisher information of time is given by *ds*^2^*/dt*^2^ = ∑_*i*_ *P*_*i*_*(d* ln *P*_*i*_*/dt)*^2^ (nats^2^*/*min^2^) = ∑_*i*_ *P*_*i*_*(d* log_2_ *P*_*i*_*/dt*^2^) (bits^2^ */*min^2^).

The information-geometric quantities 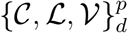 also differentiate these conditions (Fig. 3 and table S2). Firstly, we discuss the histogram of the action C. The mean value of the action 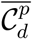 becomes larger as cell density increases 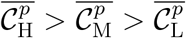 for both *p* = a and *p* = i. It reflects that the speed under the high density appeared faster than the low density. The difference of 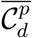 between the high and low densities is at least twice 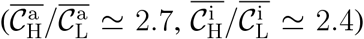. Compared with the activation and inactivation processes, 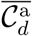 is approximately twice as much as 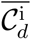. The distributions for the activation have longer tails than the inactivation. Because the action is a measure of the thermodynamic cost, these results suggest that the activation’s thermodynamic cost is larger than the inactivation’s one and becomes larger as cell density increases. Secondly, we discuss the histogram of the length ℒ. While the length does not distinguish the activation process from the inactivation process, the mean value of the length 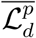 differentiates the cell densities. It reflects the fact that the peak of the spike becomes higher as the cell density increases. Finally, we discuss the histogram of the mean velocity 𝒱. The mean value of the mean velocity for the activation 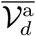 is larger than the inactivation 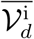. It shows that the activation speed is faster than the inactivation, which reflects the regulation speed of cell proliferation.

**Fig. 3.**
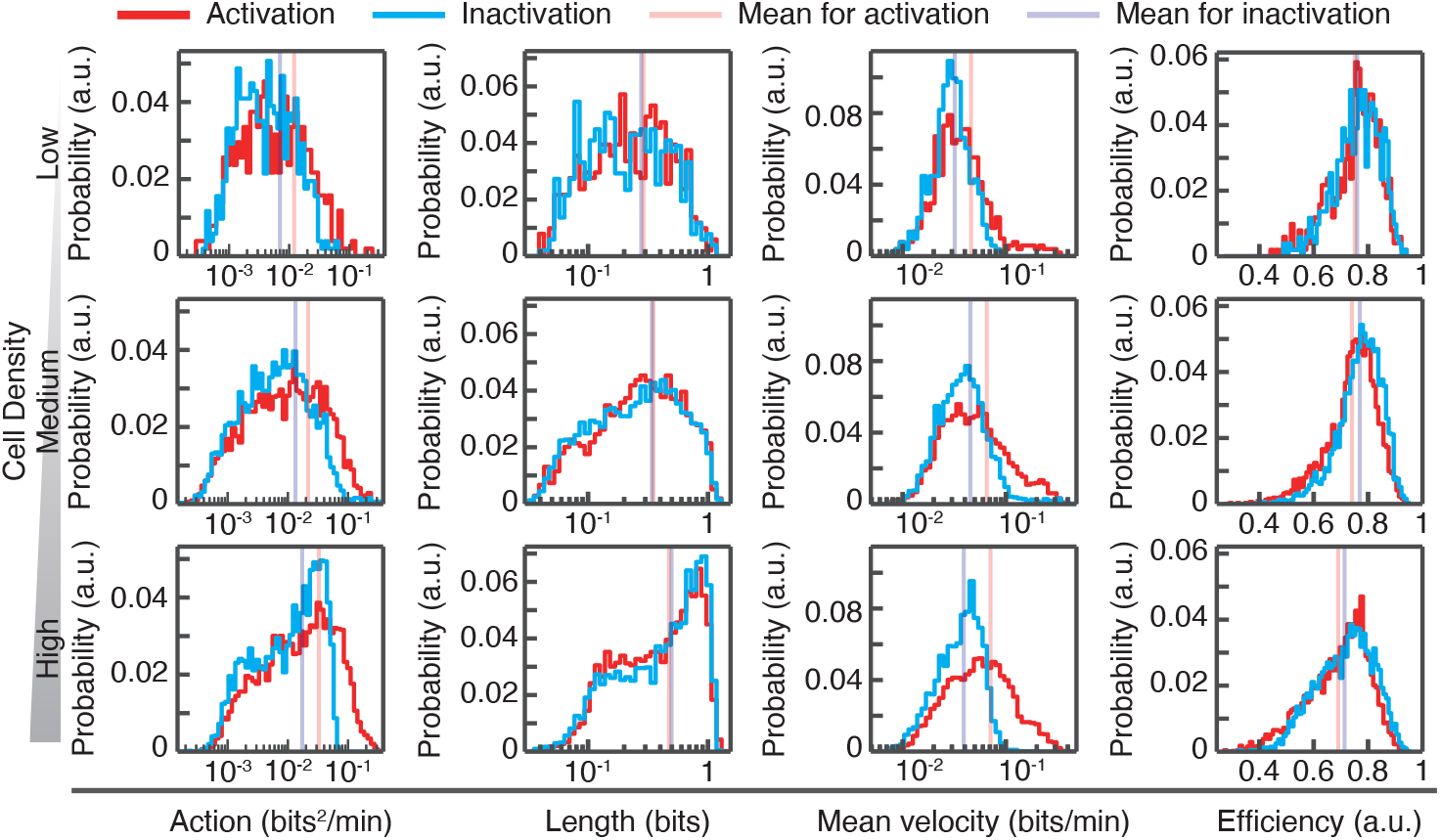
The histograms of the action 𝒞, length ℒ, mean velocity 𝒱, and efficiency *η* under the conditions of different cell densities. The sample size, mean values, variances, and results of statistical tests are listed in table S1, S2.

From these information-geometric quantities 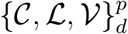, we obtained the speed limit’s efficiency 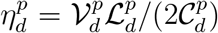 (Fig. 3 right column). The mean value of the speed limit’s efficiency is almost 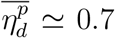 under these conditions. The distribution of 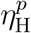 shifted to lower than those of 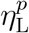 and 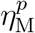 (see also table S2). The efficiencies range between 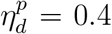 and 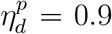, and the distributions’ shapes are biased to the higher efficiency. We can detect that the activation process is less efficient than the inactivation process 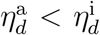 . We also find that the efficiency becomes worse in the higher cell density 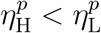 . These results indicate that the process is accelerated under the higher cell density, and the activation process is also more accelerated than the inactivation process by the thermodynamic cost. This idea is supported by the histogram of |*d/dt(ds*^2^*/dt*^2^)| (fig. S1), which becomes zero when *η* = 1. The mean values of |*d/dt(ds*^2^*/dt*^2^)| for the activation and the high cell density are larger than that for the inactivation and the low cell density, respectively (table S2). The scatter plot (fig. S2, see also table S4) suggests no correlation between 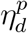 and 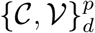, while 𝒱 and 𝒞 seem to correlate. Because cell density modulates the ERK activation’s excitability through both changes in basal and peak levels of the ERK phosphorylation by interaction with neighboring cells *(31)*, the cellular density could affect the efficiency of the pulsatile phosphorylation and the thermodynamic cost.

We confirm that these information-geometric quantities show the thermodynamic properties of the ERK phosphorylation by comparing the Raf inhibitor addition situation. The pulsatile dynamics of the ERK activation are generated by stochastic noise from the Raf and feedback loops *(31, 32)*, and the Gibbs free energy difference of the ERK phosphorylation is induced by stimulus *s*′ from the upstream pathway, including the Raf *(32)*. Thus, the Raf inhibitor addition affects the Gibbs free energy change of the ERK phosphorylation and these information-geometric quantities. To ensure these relationships, we measured the ERK phosphorylation dynamics under the Raf inhibitor (SB590885) addition and compared its dynamics with the original dynamics before adding the Raf inhibitor (Fig. 4). Of note, it is well-known that a low dose of the Raf inhibitor could paradoxically activate the ERK signaling through the Raf dimerization *(35)*. The condition of the cell density is the medium in this experiment. The application of a low dose (100 nM) of the Raf inhibitor immediately activated the ERK, and the ERK activity demonstrated slower dynamics than that before the Raf inhibitor treatment and after the activation *(31)* (Fig. 4A and movie S2). This result implies that the thermodynamic cost of the ERK phosphorylation is immediately increased when adding the Raf inhibitor. After the dynamics of the relaxation on the upstream pathways, the Raf inhibitor addition generally decreases the thermodynamic cost of the ERK phosphorylation.

**Fig. 4.**
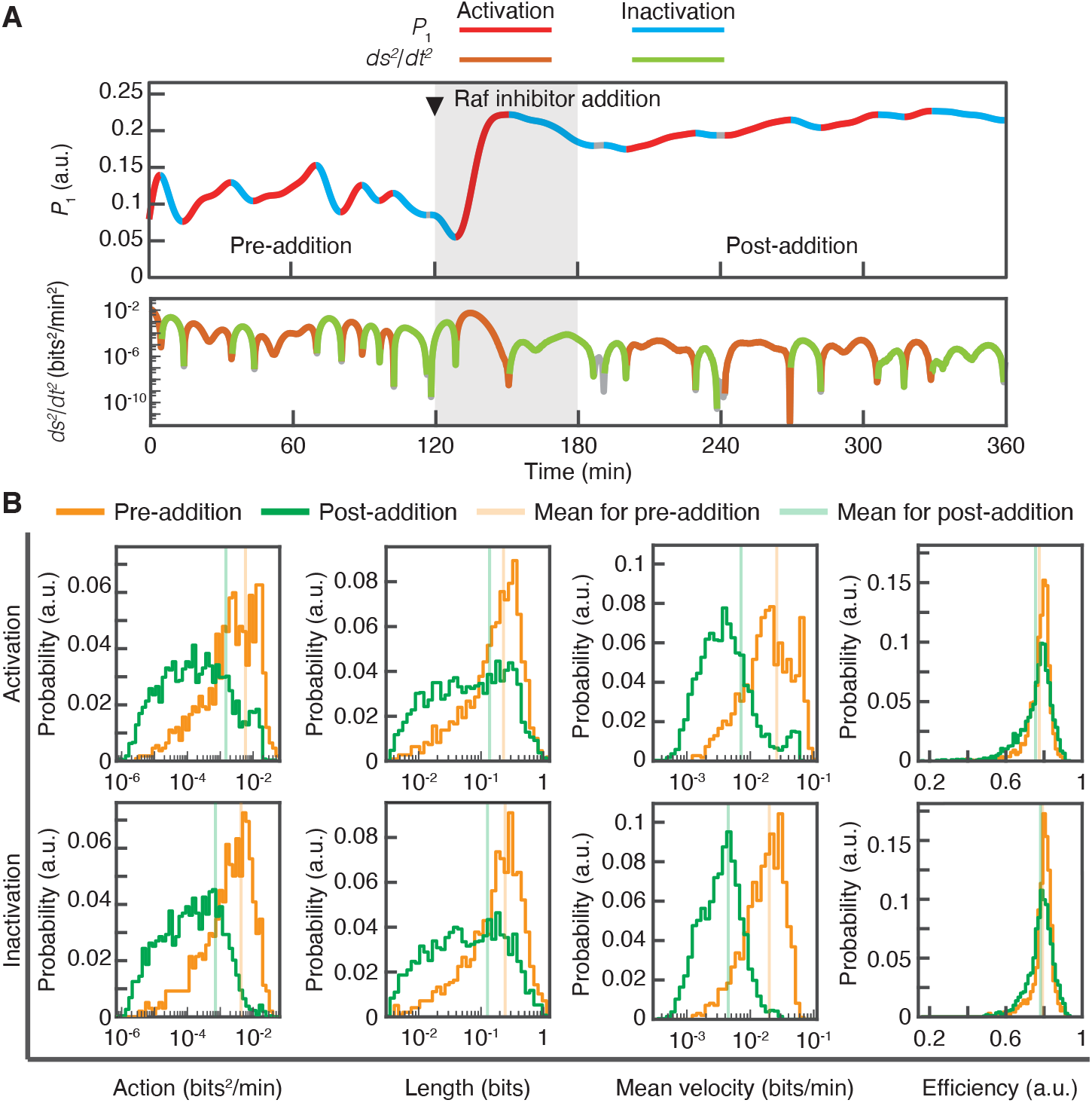
The dynamics of the ERK phosphorylation under the Raf inhibitor addition. (**A**) Representative dynamics under the Raf inhibitor addition. The Raf inhibitor (100 nM of SB590885) was applied 120 min after the imaging initiation. We set the pre-addition region as the period from 0 min to 120 min, and the post-addition region as the period after 180 min. (**B**) The histograms of the action 𝒞, length ℒ, mean velocity 𝒱, and the efficiency *η* before (pre-addition) and after (post-addition) adding the Raf inhibitor. The sample size, mean values, variances, and results of statistical tests are shown in table S1, S2.

In Fig. 4B, we show the histograms of these information-geometric quantities and the efficiency 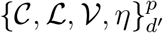 before *(d*′ = pre) and after *(d*′ = post) the Raf inhibitor addition, where the subscript *d*′ ∈ {pre, post} regards the Raf inhibitor addition. The mean value of the action 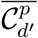 decreases when the Raf is inhibited 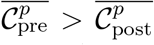, and the mean values differ by one order of magnitude (see also table S3). This result shows that the Raf inhibitor addition reduces the thermodynamic cost of the ERK phosphorylation. The mean velocity and length also decrease when the Raf inhibitor is added, 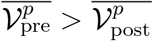 and 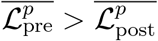, while the transition time *t*_1_ − *t*_0_ becomes longer by the Raf inhibitor addition (fig. S3 and see also table S3). When adding the Raf inhibitor, cell proliferation is promoted because the speed of the regulators such as the ELK1 and RSK becomes slow *(35)*. Thus, the change of the mean velocity explains the behavior of cell proliferation under the Raf inhibitor addition. Surprisingly, the mean value of the efficiency is robust to the Raf inhibitor addition 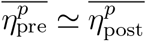, while the peak of the histogram is decreased after adding the Raf inhibitor. Moreover, the efficiency 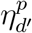 seems not to correlate with 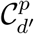 nor 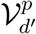 (fig. S4 and see also table S4), and this robustness would not come from an artifact correlation between information-geometric quantities. The efficiency compensation of the ERK phosphorylation might exist when the upstream pathways are perturbed by the inhibitor.

In summary, we introduced information-geometric quantities as thermodynamic measures and evaluated the speed limit’s efficiency for the ERK phosphorylation dynamics in living cells at the single-cell level. Our method quantitatively clarifies the amount of information transferred in living systems based on information geometry and the conversion efficiency from the thermodynamic cost to the intrinsic speed, which complements other studies about measurements of informational quantities in biological systems such as the mutual information in signal transduction *(36,37)* and the Fisher information matrix on molecular networks *(38)*. Our method is generally applicable to the activation process, and it widely exists in signaling pathways such as the G-protein coupled receptor pathways for sensation and the RTK signaling pathways for cell proliferation. For example, the state change from the active state *A*_on_ to the inactive state *A*_off_ in the G-protein coupled receptor corresponds to its conformational change by small re-arrangements accompanying the ligand-binding. It is interesting to evaluate the speed limit’s efficiency for other activation processes on an equal footing with this ERK phosphorylation in living NRK-52E cells. Our approach has great potential for other biological applications, which might clarify an efficient signal transduction mechanism and lead to novel insights into living systems as an information processing unit driven by the thermodynamic cost.

## Acknowledgements

S. I. and Keita Ashida thank Kohei Yoshimura for discussions of chemical thermodynamics, S. I thanks Keita Kamino for the discussions of experimental difficulties in measurement of information quantity of living cells, and Kazuhiro Aoki thanks Yohei Kondo for helpful discussions of thermodynamics. S. I. also thanks Kiyoshi Kanazawa for careful reading of this manuscript.

## Author contributions

S. I. proposed the main method based on information geometry and designed the research. Kazuhiro Aoki performed experiments. Keita Ashida analyzed the data. All authors discussed the analytical results of the experiment and wrote the paper.

## Data and materials availability

The data sets generated and analyzed during the current study are available from the corresponding author on reasonable request. The source codes used in the current study are available from the corresponding author on reasonable request.

## Supplementary Materials for

### Materials and Methods

#### Measurement of the phosphorylated ERK fraction

We measured the ERK phosphorylation, as previously described *(39)*. The EKAREV-NLS stable-expression NRK-52E cell lines (NRK-52E/ERKAREV-NLS cells) *(31)* were used, and the EKAREV-NLS is the genetically encoded ERK sensor used in Komatsu *et al*., 2011 *(34)*.

The NRK-52E/EKAREV-NLS cells were maintained with the Dulbecco’s Modified Eagle Medium (DMEM; ThermoFisher), 10 % Fetal bovine serum (FBS; Sigma), and 10 µg/mL Blasticidin S (Invitrogen). The cells were seeded at a specific concentration (Low: 2.0 × 10^3^ cells/cm^2^, Medium: 2.0 ×10^4^ cells/cm^2^, High: 2.0 × 10^5^ cells/cm^2^) on glass-bottom dishes (IWAKI). One day after the seeding, the time-lapse imaging was performed. The culture media was replaced with the FluoroBrite (ThermoFisher), 5 % FBS (Sigma), and 1 × Glutamax (ThermoFisher) 3–6 hours before starting the time-lapse imaging. We used an inverted microscope (IX81; Olympus) equipped with a CCD camera (CoolSNAP K4; Roper Scientific) and an excitation light source (Spectra-X light engine; Lumenncor). Optical filters were as follows: an FF01-438/24 excitation filter (Semrock), an XF2034 (455DRLP) dichroic mirror (Omega Optical), and two emission filters (FF01-483/32 for CFP and FF01-542/27 for YFP (Semrock)). Images were acquired every 20 sec (the exposure time was 100 ms) with binning 8 × 8 on MetaMorph software (Universal Imaging) with an IX2-ZDC laser-based autofocusing system (Olympus). A ×20 lens (UPLSAPO 20X; Olympus, numerical aperture: 0.75) was used. The size of the field of view is 757.76 × 757.76 µm. The temperature and CO_2_ concentration were maintained at 37°C and 5 % during the imaging with a stage-top incubator (Tokai hit). For the Raf inhibitor experiment, the experiment was performed under the same condition as the medium cell density condition. We applied SB590885 (Selleck Chemicals) (a final concentration is 100 nM) 2 hours after the imaging initiation. The numbers of trials for each experimental condition are two.

We used the same relation between the FRET/CFP ratios of the EKAREV-NLS and the phosphorylated ERK fraction (pTpY-ERK2) from the western blotting in Fig. 2B, as described previously *(31)*. In brief, the phosphorylated ERK fraction was quantified by the Phos-tag western blotting *(40)* in HeLa cells stimulated with different concentrations of 12-O-Tetradecanoylphorbol 13-acetate (TPA; Sigma) for 30 min to induce the ERK phosphorylation. Under the same condition, HeLa cells stably expressing EKAREV-NLS were imaged, followed by quantifying the average FRET/CFP ratios. Finally, the FRET/CFP ratios were plotted as a function of the phosphorylated ERK fraction with the fitted curve of the Hill equation shown in Fig. 2B by the Solver Add-in in Excel (Microsoft).

#### Data analysis

For the imaging analysis, Fiji was used. The background was subtracted by the Subtract Background Tool, and after that, the nuclei were tracked with a custom-made tracking program. Only the cells tracked over the entire images were used to calculate the information-geometric quantities. The phosphorylated ERK fraction was calculated based on the FRET/CFP ratios with the Hill equation shown in Fig. 2B.

For analysis of the time-series phosphorylated ERK fractions, MATLAB 2019b (MathWorks) was used. The low-pass filter whose cutoff frequency is 0.005 Hz was used for the cell density experiment data with the designfilt function in the Signal Processing Toolbox to reduce the noise. The low-pass filter whose cutoff frequency is 1*/*1200 Hz was used for the Raf inhibitor addition experiment data to detect the slower dynamics after the Raf inhibitor addition. We identified the activation and inactivation processes from the signs of the first and second derivatives. The first derivatives were numerically calculated using two data points of the phosphorylated fraction, and the second derivatives were numerically calculated using two data points of the first derivatives. We only calculated the information-geometric quantities when the phosphorylated ERK fraction difference between maximum and minimum of the process over 0.01 for the density-experiment data and 0.001 for the Raf inhibitor addition experiment data. The square of an intrinsic speed *ds*^2^*/dt*^2^ was numerically calculated using two data points of the time series phosphorylated ERK fraction. The action 𝒞, length ℒ, mean velocity 𝒱, and efficiency *η* were calculated using *ds*^2^*/dt*^2^ according to the definitions. We used the set of time series for each cell in two trials for each experimental condition to make histograms.

For the statistical analysis, R (version 3.6.3; R project) was used. The Brunner-Munzel test was performed with the brunnermunzel.test function in the brunnermunzel library (version 1.4.1). The two-sample Kolmogorov–Smirnov test was performed with the ks.test function in the stats library (version 3.6.3). The Pearson correlation coefficients were calculated and tested by the cor.test function in the stats library (version 3.6.3). The Holm method was used to control the family-wise error rates with the p.adjust function in the stats library (version 3.6.3). The sample numbers are shown in table S1.

### Supplementary Text

#### Chemical thermodynamics and information geometry for switching dynamics between the active and inactive states

This supplementary text briefly discusses a relationship between chemical thermodynamics and information geometry for switching dynamics between the active state *A*_on_ and the inactive state *A*_off_ . Let us consider the chemical reaction for the active and inactive states with stimulus *s*′,

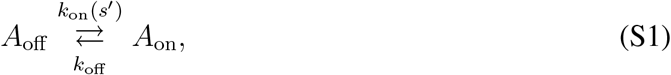

where *k*_on_*(s*′) and *k*_off_ are rate constants. Let [*A*_off_] and [*A*_on_] be concentrations of the states *A*_off_ and *A*_on_, respectively. These concentrations are nonnegative, i.e., [*A*_off_] ≥ 0 and [*A*_on_] ≥ 0. In our study, *A*_on_ corresponds to the phosphorylated ERK state, *A*_off_ corresponds to the nonphosphorylated ERK state, and *s*′ corresponds to upstream proteins’ stimulus in the Ras-Raf-MEK-ERK pathway.

At first, we consider switching dynamics between the active and inactive states. The following rate equation describes the time evolution of concentrations [*A*_off_] and [*A*_on_],

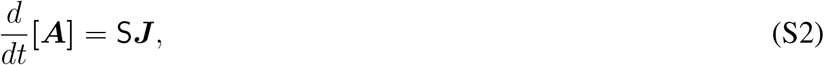

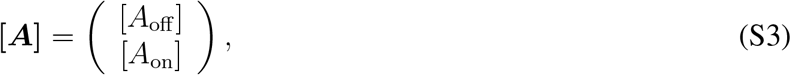

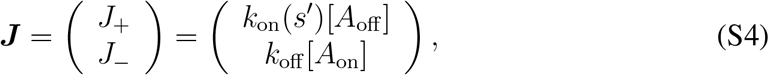

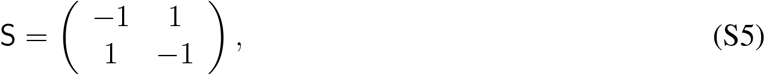

where *t* is time, the vector ***J*** implies the reaction rate, and S is called the stoichiometric matrix. Because the vector

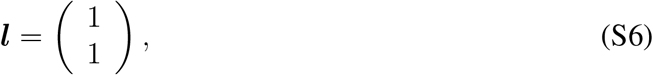

is in the left null space of S such that

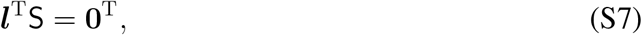

the sum of concentrations [*A*_tot_] = [*A*_on_] + [*A*_off_] is conserved

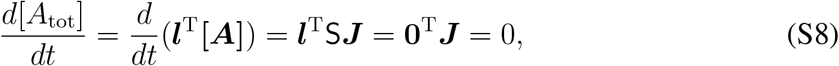

where **0** be the zero vector in two dimensions, and the symbol ^T^ implies the transpose of a vector. This conservation law *d*[*A*_tot_]*/dt* = 0 corresponds to the conservation of the total mass. If we introduce the concentration fractions

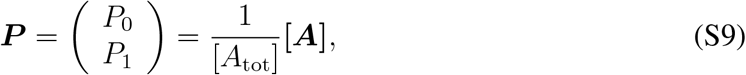

we have *P*_0_ ≥ 0, *P*_1_ ≥0, and *P*_0_ + *P*_1_ = 1. Then, *(P*_0_, *P*_1_) can be regarded as the probability distribution. Its dynamics are described by the master equation, which is equivalent to the rate equation Eq. (S2),

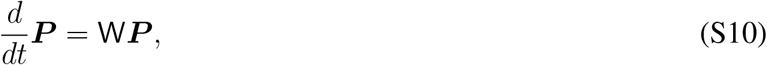

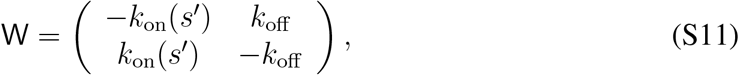

where W is called the rate matrix. The equilibrium distribution is given by

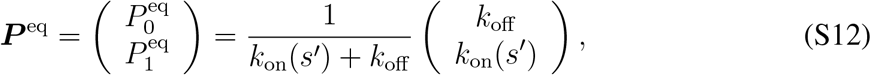

which satisfies the detailed balance condition

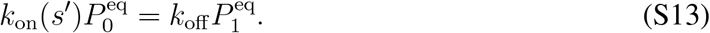

This equilibrium distribution ***P*** ^eq^ generally depends on the stimulus *s*′. The concentrations at equilibrium corresponding to ***P*** ^eq^ are given by

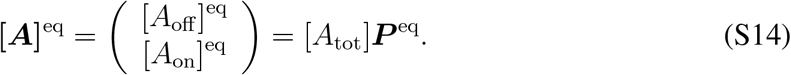

The ratio of two rate constants *k*_on_*(s*′)*/k*_off_ implies a thermodynamic property of the system. This fact is known as the local detailed balance condition such that

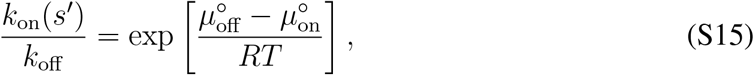

where *R* is the gas constant, *T* is the temperature of the solvent, and 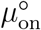 and 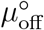 are the standard chemical potentials corresponding to the active and inactive states. We now define the chemical potentials *µ*_on_([*A*_on_]) and *µ*_off_ ([*A*_off_]) as

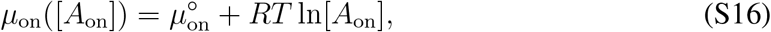

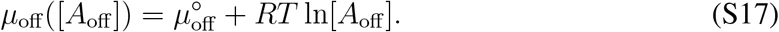

The local detailed balance condition Eq. (S15) can be regarded as the equivalence of the chemical potential at equilibrium

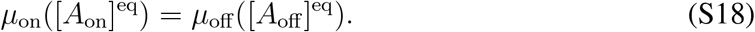

If we define the vector of the chemical potential at equilibrium as

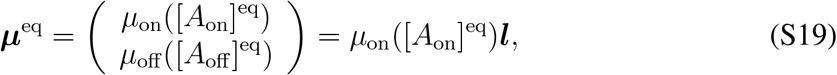

this vector is in the left null space of S, i.e., ***µ***^eqT^S = **0**^T^. Then, the quantity ***µ***^eqT^**[*A*]** is conserved,

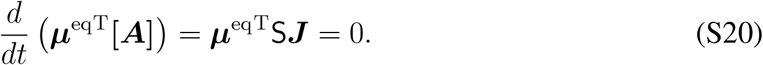

Next, we discuss the chemical thermodynamics of a dilute solution with the temperature *T* kept constant. The Gibbs free energy per unit volume *G*(**[*A*]**) is defined as

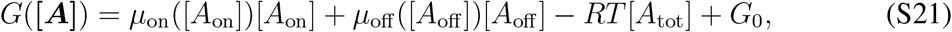

where *G*_0_ is a constant. This Gibbs free energy *G*(**[*A*]**) satisfies the relations in chemical thermodynamics,

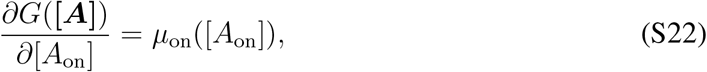

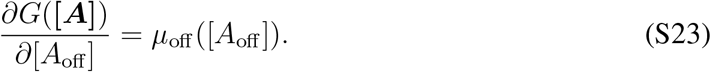

From Eq. (S20), we obtain

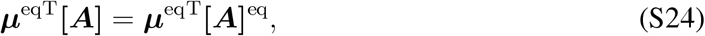

and therefore the Gibbs free energy at equilibrium *G*(**[*A*]**^eq^) is expressed as

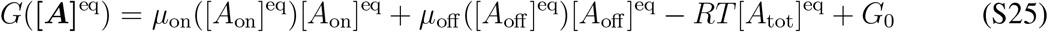

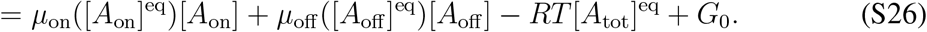

The Gibbs free energy difference *G*(**[*A*]**) − *G*(**[*A*]**^eq^) is given by

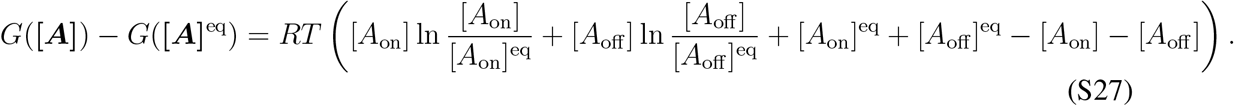

If we use the definition of the *f* -divergence

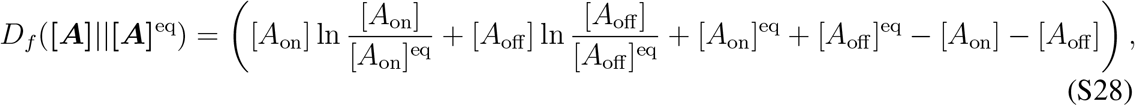

which is the generalization of the Kullback-Leibler divergence for the positive measure space, the Gibbs free energy difference is given by

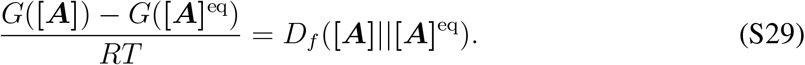

From the nonnegativity of the *f* -divergence *D*_*f*_ ([*A*]||[*A*]^eq^) ≥ 0, we obtain

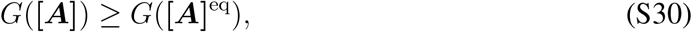

which implies that the Gibbs free energy at equilibrium takes the smallest value. If we consider the conservation of the total mass [*A*_on_]^eq^ +[*A*_on_]^eq^ = [*A*_on_]+[*A*_on_] and introduce the probability distributions ***P*** and ***P*** ^eq^, the *f* -divergence is rewritten as the Kullback-Leibler divergence,

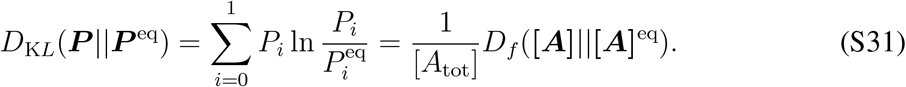

The time derivative of the Gibbs free energy is also given by the *f* -divergences

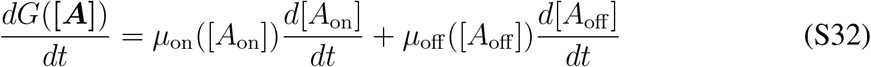

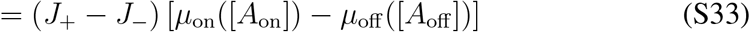

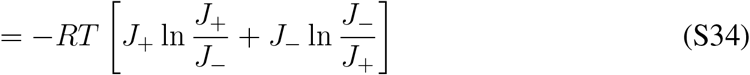

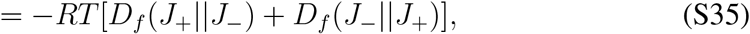

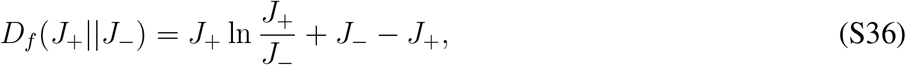

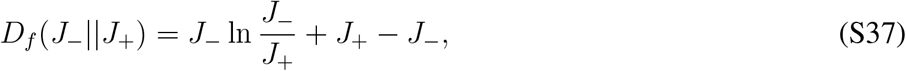

where we used the local detailed balance condition Eq. (S15), which is equivalent to *µ*_on_([*A*_on_])− *µ*_off_ ([*A*_off_]) = *RT* ln*(J*_−_*/J*_+_). From the nonnegativity of the *f* -divergence, we obtain

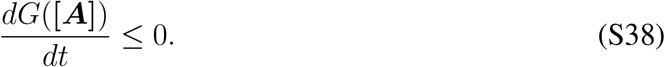

The entropy production *σ*(**[*A*]**) is defined as

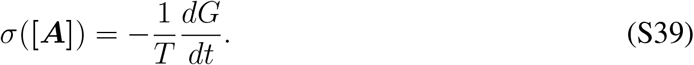

Therefore, *dG*(**[*A*]**)*/dt* ≤ 0 can be regarded as the second law of thermodynamics

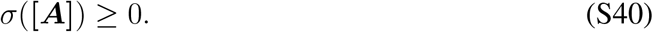

We here introduce information geometry. In information geometry, the square of the line element *ds*^2^ is given by the second-order Taylor expansion of the Kullback-Leibler divergence

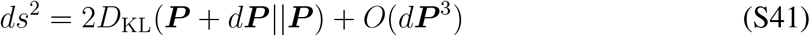

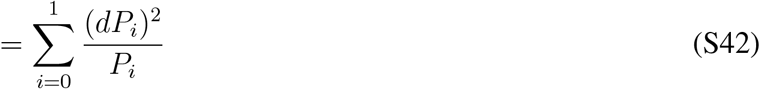

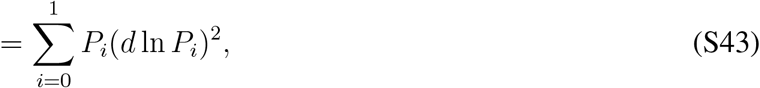

where *d****P*** is an infinitesimal difference of two probabilities, which satisfies ∑_*i*_ *dP*_*i*_ = 0. The Fisher information of time is also defined as

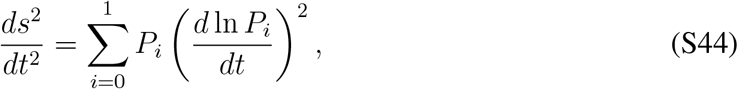

and its square root implies the intrinsic speed 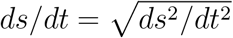 in information geometry. We can also introduce information geometry for chemical reaction networks. The square of line element for chemical reaction networks 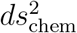 is given by the second-order Taylor expansion of the *f* -divergence

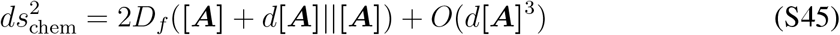

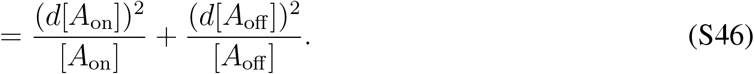

We obtain the relation between two geometries *ds*^2^ and 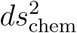 as

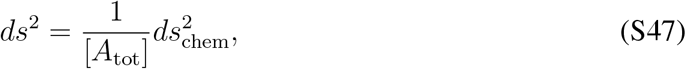

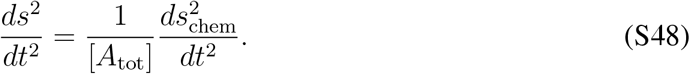

Because the *f* -divergence is related to the Gibbs free energy, we can consider a thermodynamic interpretation of information geometry. Under near-equilibrium conditions **[*A*]** = **[*A*]**^eq^ +*d***[*A*]**, we have

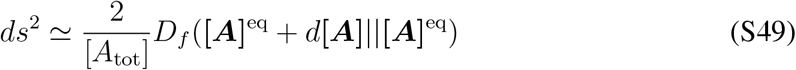

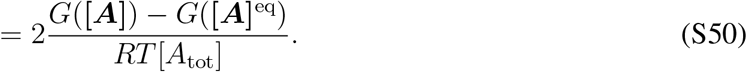

Therefore, *ds*^2^ can be interpreted as the Gibbs free energy difference under near-equilibrium conditions. We also show that the Fisher information of time *ds*^2^*/dt*^2^ is given by

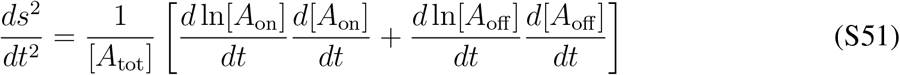

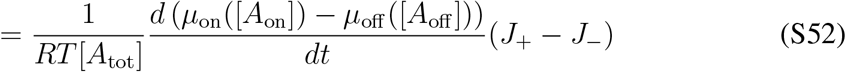

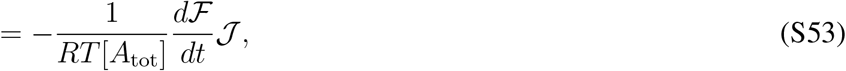

where the flow is defined as 𝒥 = *J*_+_ − *J*_−_ and the force is defined as ℱ = *µ*_off_ ([*A*_off_]) − *µ*_on_([*A*_on_]) = *RT* ln*(J*_+_*/J*_−_). In this notation, the entropy production rate is given by *σ* = ℱ𝒥 */T* . In terms of the entropy production rate, the Fisher information of time is given by

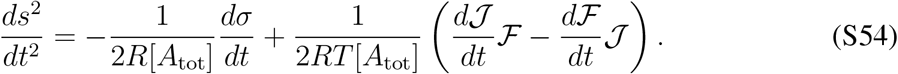

Under near-equilibrium conditions *J*_+_ ≃ *J*_−_, we have 𝒥 ≃ *Lℱ* with the Onsager’s coefficient *L* = *RT/(k*_off_ [*A*_on_]^eq^) and

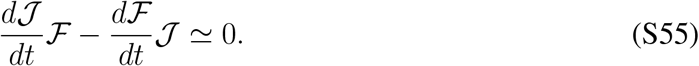

Thus, the Fisher information of time gives the time derivative of the entropy production rate under near-equilibrium conditions

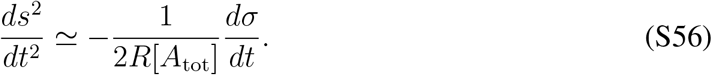

We also discuss the thermodynamic meaning of the action for a transition from time *t* = *t*_0_ to *t* = *t*_1_, defined as

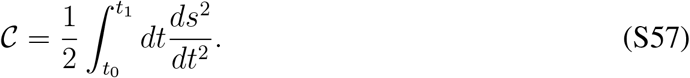

Under near-equilibrium conditions, we obtain

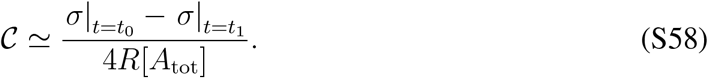

In a relaxation process, the system can be in equilibrium at the final time *t* = *t*_1_, and the entropy production rate vanishes at the final time 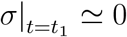. Therefore, the action can be proportional to the entropy production rate at the initial time *t* = *t*_0_,

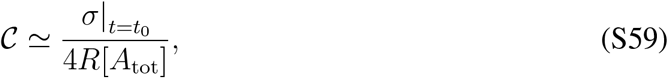

if the system is under near-equilibrium conditions and reach equilibrium at the final time *t* = *t*_1_. That is why we regard the action 𝒞 as a thermodynamic cost. In an experiment of the FRET measurement, we can easily estimate the Fisher information of time *ds*^2^*/dt*^2^ because it only consists of measurable quantities, i.e., the nonphosphorylated and phosphorylated ERK fractions *P*_0_ and *P*_1_ and their change speed *dP*_0_*/dt* and *dP*_1_*/dt*. On the other hand, it is hard to measure the Gibbs free energy *G* itself because we need prior knowledge about concentrations at equilibrium [*A*_on_]^eq^ and [*A*_off_]^eq^ to estimate *G*. It is also hard to measure the entropy production rate *σ* because we need prior knowledge about the rate constants *k*_on_*(s*′) and *k*_off_ to estimate *σ* via *J*_+_ and *J*_−_.

Information theoretically, we can discuss the meaning of *ds*^2^*/dt*^2^ even for a system far from equilibrium. We here introduce the observable *(f*_0_, *f*_1_) corresponding to *(P*_0_, *P*_1_), and its ensemble average is defined as ⟨*f* ⟩_*t*_ = *P*_0_*(t)f*_0_ + *P*_1_*(t)f*_1_. The square of the time derivative of ⟨*f* ⟩_*t*_ is given by

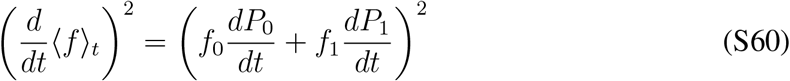

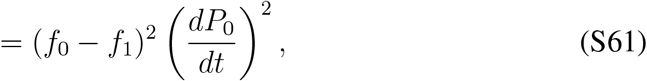

which implies the time-response of the observable. The variance of the observable is given by

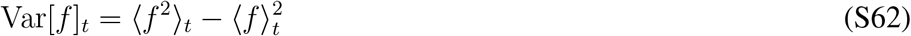

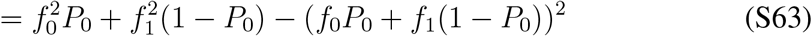

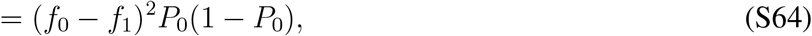

which is the fluctuation of the observable. Thus, the fluctuation-response ratio is calculated as

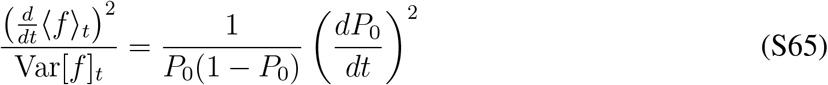

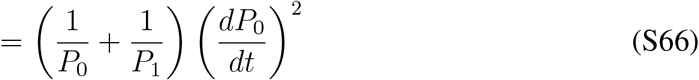

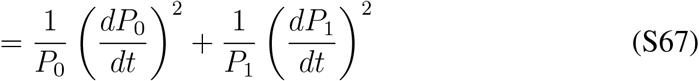

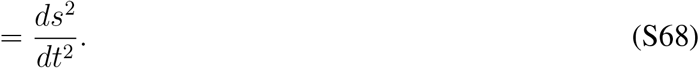

Therefore, the Fisher information of time implies the fluctuation-response ratio even for a system far from equilibrium.

This relation between the fluctuation-response ratio and the Fisher information of time is also generalized for the time derivative of the Gibbs free energy. The square of the entropy production is calculated as

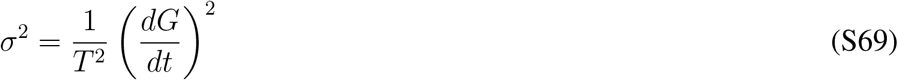

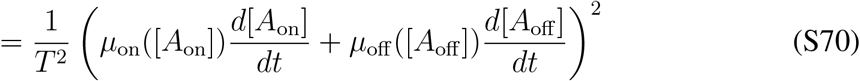

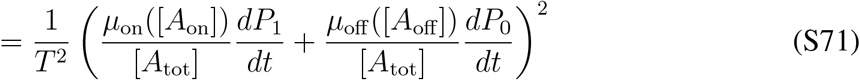

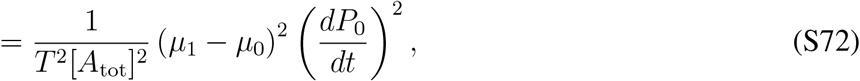

and the fluctuation of the chemical potential is calculated as

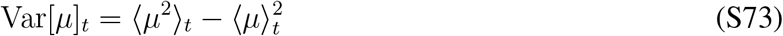

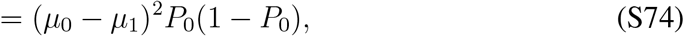

where we used the notation *µ*_0_ = *µ*_off_ ([*A*_off_]) and *µ*_1_ = *µ*_on_([*A*_on_]). Thus, we obtain the relation between the entropy production rate and the Fisher information of time,

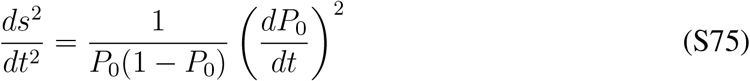

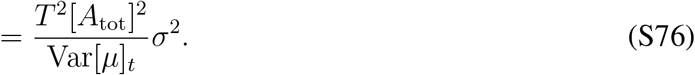

Thus, the Fisher information of time is proportional to the square of the entropy production rate.

We finally discuss the length ℒ defined as

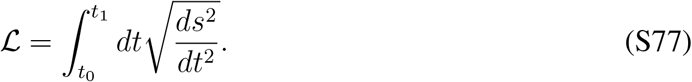

We here introduce the change of variables 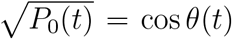 and 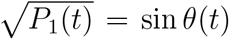, which satisfies the normalization of the probability *P*_0_*(t)* + *P*_1_*(t)* = 1. Then, the Fisher information of time is given by

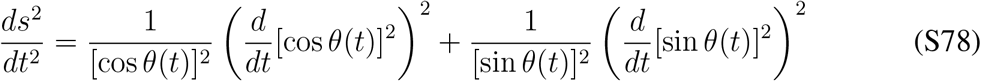

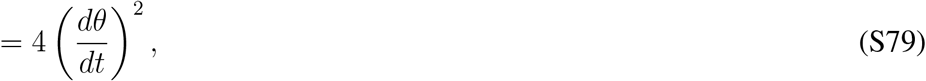

which implies that *ds*^2^ provides a differential geometry of the circle of radius 2. Thus, the length is calculated as

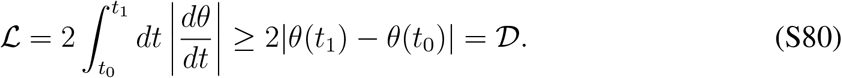

The lower bound 𝒟 is the geodesic length on the circle of radius 2. If we assume that the time evolution of *P*_1_ from time *t*_0_ to *t*_1_ is monotonic, the equality holds ℒ = 𝒟. In the main text, we consider ℒ= 𝒟 for the activation and inactivation processes because *P*_1_ in the activation (inactivation) process is monotonically increasing (decreasing) in time. From the identity

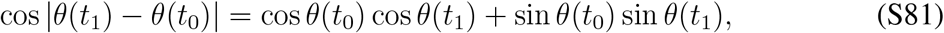

the length can be calculated as

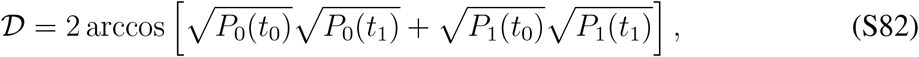

where the angle 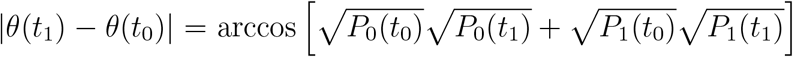 is known as the Bhattacharyya angle.

**Fig. S1.**
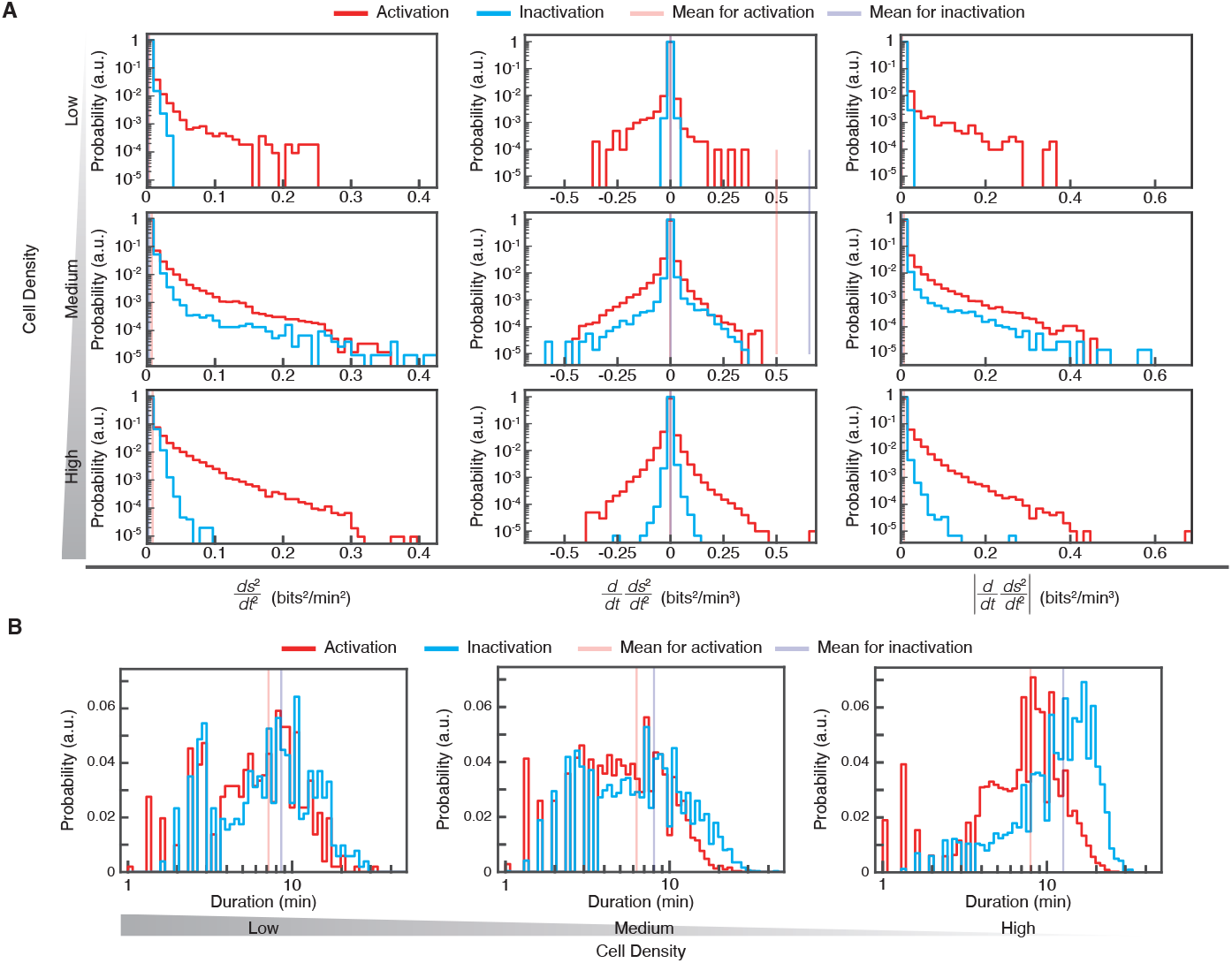
The histograms of the square of the intrinsic speed and related quantities at the low, medium, and high cell densities. (**A**) The histograms of *ds*^2^*/dt*^2^, *d/dt(ds*^2^*/dt*^2^), and |*d/dt(ds*^2^*/dt*^2^)| at each cell density. (**B**) The histograms of the duration *(t*_1_ − *t*_0_) of the activation and inactivation processes under each cell density. The mean values, variances, and results of statistical tests are listed in table S2.

**Fig. S2.**
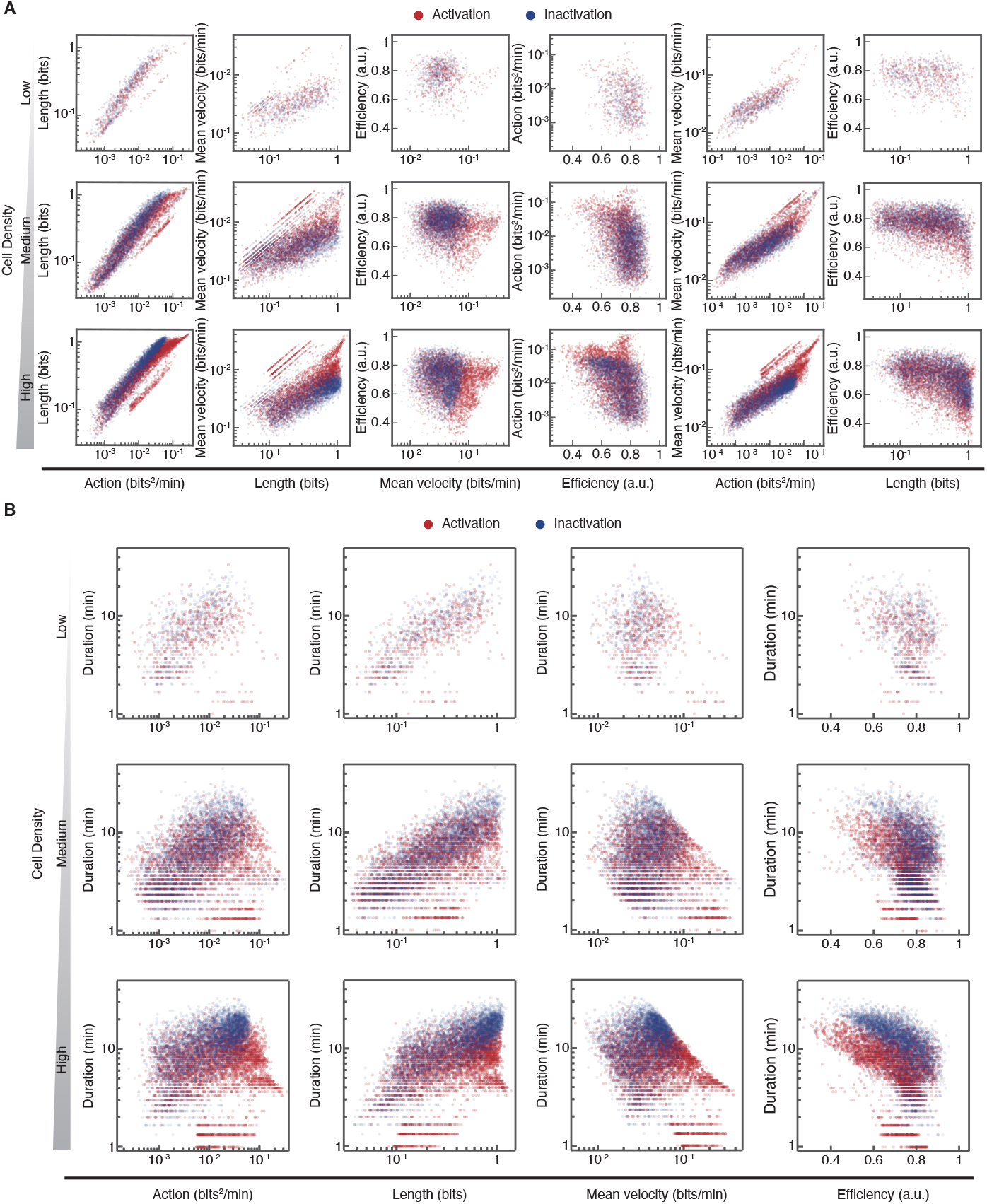
The correlation between two information-geometric quantities at each cell density. (**A**) The scatter plots of the action 𝒞, the length ℒ, the mean velocity 𝒱, and the efficiency *η*. (**B**) The scatter plots of the duration *t*_1_ − *t*_0_ versus information-geometric quantity (𝒞, ℒ, 𝒱, *η)*. The Pearson correlation coefficients and results of statistical tests are listed in table S4.

**Fig. S3.**
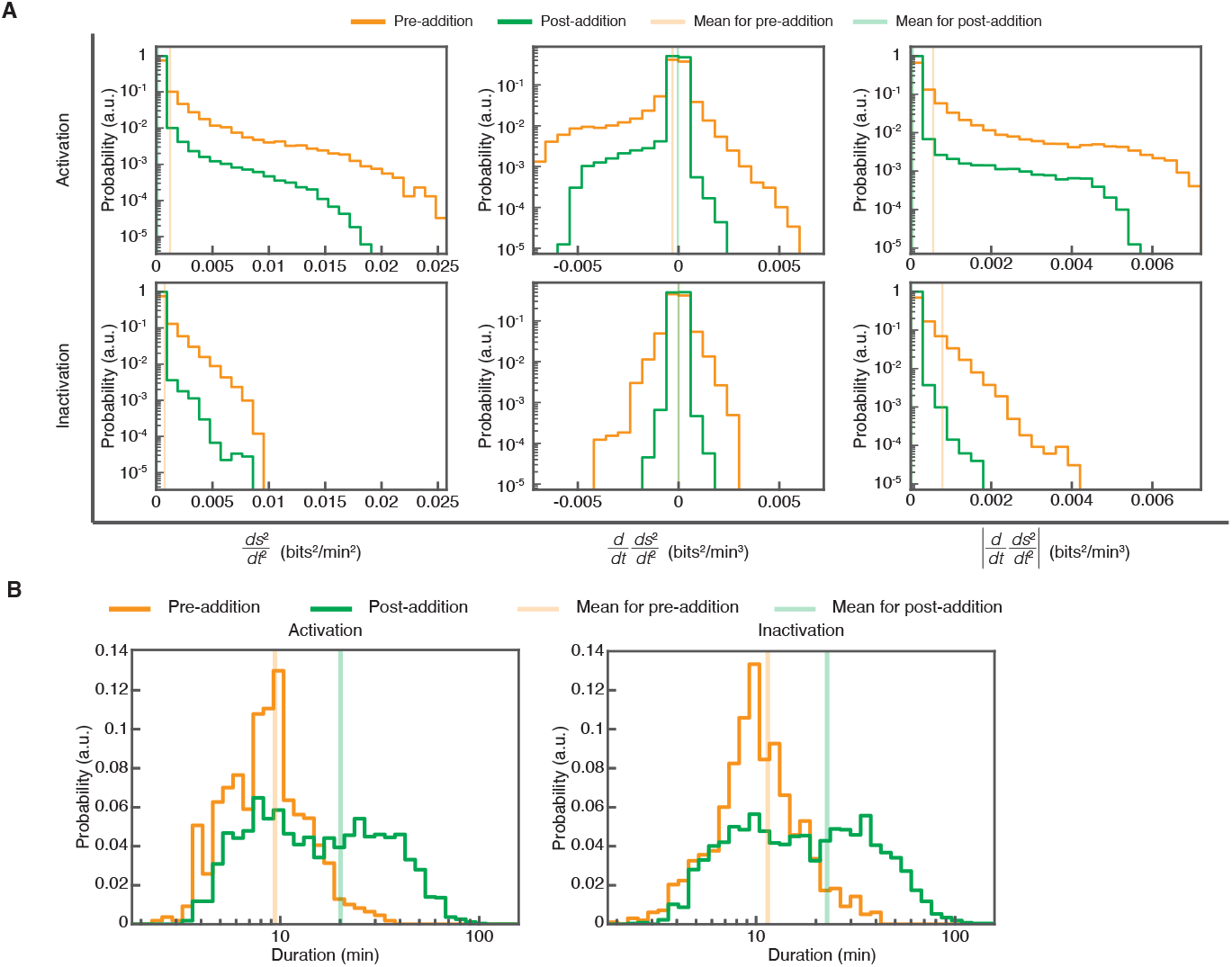
The histograms of the square of the intrinsic speed and related quantities before and after the Raf inhibitor addition. (**A**) The histograms of *ds*^2^*/dt*^2^, *d/dt(ds*^2^*/dt*^2^), and |*d/dt(ds*^2^*/dt*^2^)| of the activation and inactivation processes before (pre-addition) and after (post-addition) the Raf inhibitor addition. (**B**) The histograms of the duration *t*_1_ − *t*_0_ of the activation and inactivation process before (pre-addition) and after (post-addition) the Raf inhibitor addition. The mean values, variances, and results of statistical tests are listed in table S3.

**Fig. S4.**
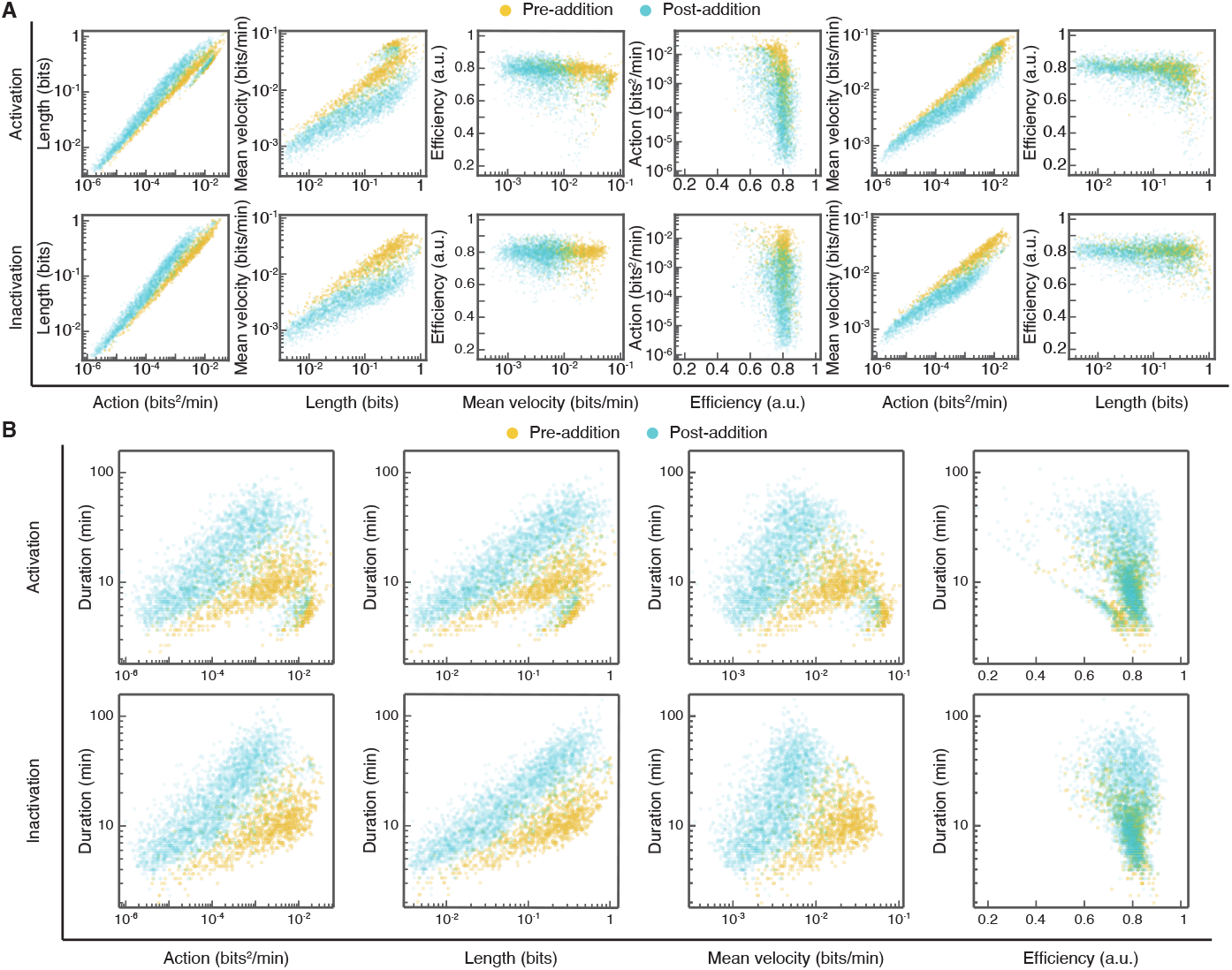
The correlation between two information-geometric quantities before and after the Raf inhibitor addition. (**A**) The scatter plots of the action 𝒞, the length ℒ, the mean velocity 𝒱, and the efficiency *η* before (pre-addition) and after (post-addition) the Raf inhibitor addition. (**B**) The scatter plots of the duration *t*_1_ − *t*_0_ versus information-geometric quantity (𝒞, ℒ, 𝒱, *η)* before (pre-addition) and after (post-addition) the Raf inhibitor addition. The Pearson correlation coefficients and results of statistical tests are listed in table S4.

**Table S1.** The sample size used in the analysis of each experiment. We listed the sample size for the experiments under the different cell densities and the experiments of the periods before and after the Raf inhibitor addition.

| Condition | Number of cells | Number of activation | Number of inactivation |
| --- | --- | --- | --- |
| Low density (L) | 31 | 507 | 513 |
| Medium density (M) | 306 | 3,130 | 3,131 |
| High density (H) | 755 | 4,425 | 4,029 |
| Before the Raf inhibitor addition (Pre) | 209 | 1,085 | 978 |
| After the Raf inhibitor addition (Post) | 209 | 2,716 | 2,628 |

**Table S2.**
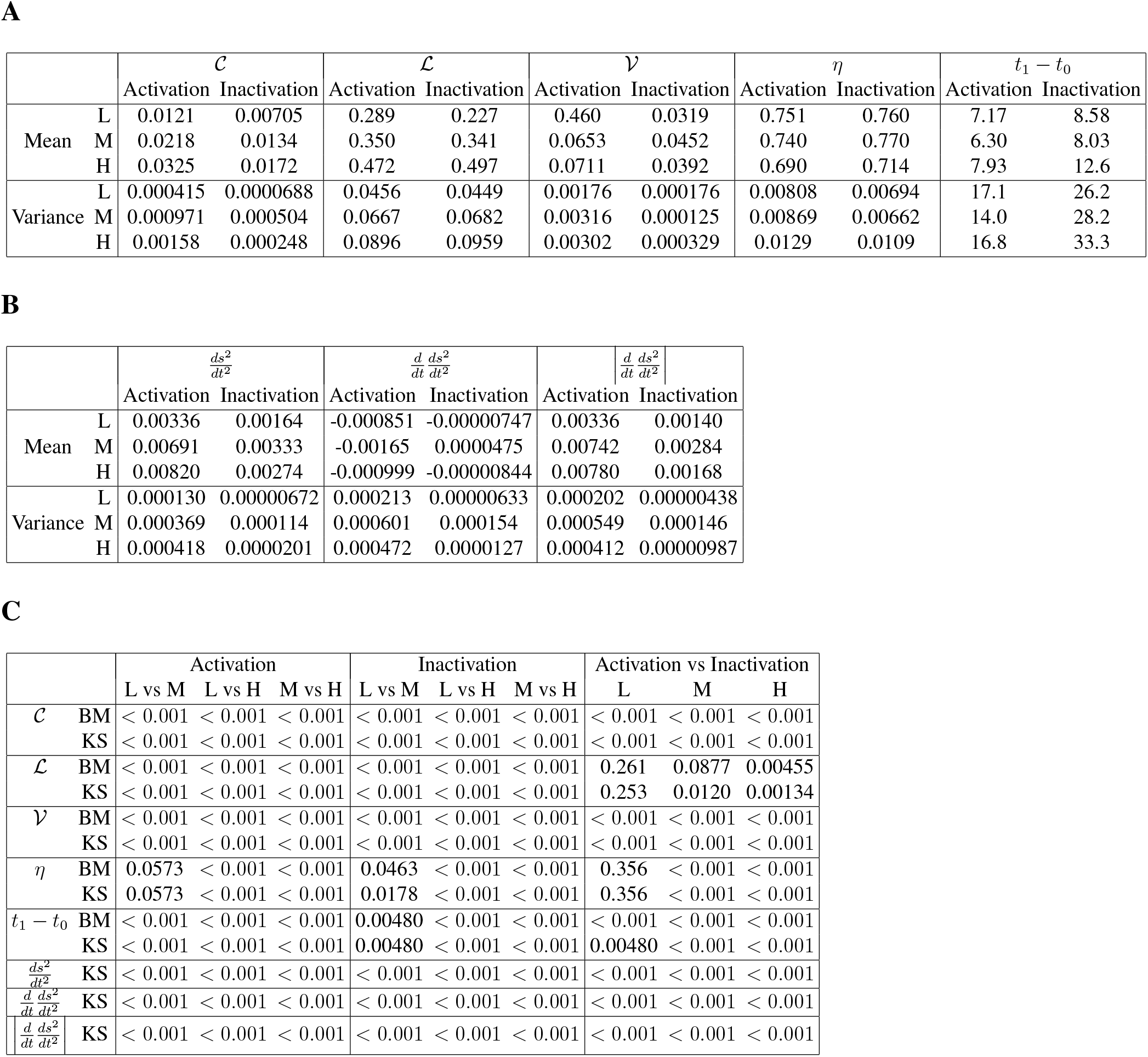
The statistics for the experiment at each cell density. (**A**) The mean values (Mean) and variances (Variance) of the action 𝒞, the length ℒ, the mean velocity 𝒱, the efficiency *η*, and the duration *t*_1_ − *t*_0_ in the activation and inactivation processes at the low (L), medium (M), and high (H) cell densities. (**B**) The mean values (Mean) and variances (Variance) of *ds*^2^*/dt*^2^, *d/dt(ds*^2^*/dt*^2^), and |*d/dt(ds*^2^*/dt*^2^)| in the activation and inactivation processes at the low (L), medium (M), and high (H) cell densities. (**C**) The p-values comparing between two cases. We used the Brunner-Munzel (BM) method and the Kolmogorov–Smirnov (KS) method. We compare two cell densities and two processes, where ‘vs’ denotes the comparison of two cases, and L, M, and H indicate the low, medium, and high cell densities. The p-values below 0.001 represent as *<* 0.001.

**Table S3.** The statistics for the experiment of the Raf inhibitor addition. (**A**) The mean values (Mean) and variances (Variance) of the action 𝒞, the length ℒ, the mean velocity 𝒱, the efficiency *η*, and the duration *t*_1_ − *t*_0_ in the activation and inactivation processes before (Pre) and after (Post) the Raf inhibitor addition. (**B**) The mean values (Mean) and variances (Variance) of *ds*^2^*/dt*^2^, *d/dt(ds*^2^*/dt*^2^), and |*d/dt(ds*^2^*/dt*^2^)| in the activation and inactivation processes before (Pre) and after (Post) the Raf inhibitor addition. **c** The p-values comparing between two cases, before (Pre) and after (Post) the Raf inhibitor addition. We used the Brunner-Munzel (BM) method and the Kolmogorov–Smirnov (KS) method. We calculated the p-values for the action 𝒞, the length ℒ, the mean velocity 𝒱, the efficiency *η*, the duration *t*_1_ − *t*_0_, *ds*^2^*/dt*^2^, *d/dt(ds*^2^*/dt*^2^), and |*d/dt(ds*^2^*/dt*^2^)|. The p-values below 0.001 represent as *<* 0.001.

| | | $\mathcal{C}$ | | $\mathcal{L}$ | | $\mathcal{V}$ | | $\eta$ | | $t_1 - t_0$ | |
| --- | --- | --- | --- | --- | --- | --- | --- | --- | --- | --- | --- |
|  |  | Activation | Inactivation | Activation | Inactivation | Activation | Inactivation | Activation | Inactivation | Activation | Inactivation |
| Mean | Pre | 0.00581 | 0.00432 | 0.230 | 0.243 | 0.0261 | 0.0199 | 0.775 | 0.792 | 9.42 | 11.5 |
|  | Post | 0.00148 | 0.000717 | 0.138 | 0.127 | 0.00712 | 0.00448 | 0.756 | 0.781 | 20.1 | 22.7 |
| Variance | Pre | 0.0000569 | 0.0000284 | 0.0305 | 0.0385 | 0.000425 | 0.000145 | 0.00394 | 0.00298 | 23.6 | 39.0 |
|  | Post | 0.0000117 | 0.00000378 | 0.0269 | 0.0247 | 0.000105 | 0.0000133 | 0.00861 | 0.00439 | 229 | 320 |

**B**
| | | $\frac{ds^2}{dt^2}$ | | $\frac{d}{dt} \frac{ds^2}{dt^2}$ | | $\left\vert \frac{d}{dt} \frac{ds^2}{dt^2} \right\vert $ | |
| --- | --- | --- | --- | --- | --- | --- | --- |
|  |  | Activation | Inactivation | Activation | Inactivation | Activation | Inactivation |
| Mean | Pre | 0.00123 | 0.000754 | -0.000297 | $-8.07 \times 10^{-8}$ | 0.000559 | 0.0000272 |
| | Post | 0.000148 | 0.0000631 | -0.0000363 | $-5.20 \times 10^{-9}$ | 0.0000535 | 0.0000133 |
| Variances | Pre | 0.00000717 | 0.00000130 | 0.00000139 | 0.000000199 | 0.00000117 | 0.000000125 |
| | post | 0.000000574 | $4.61 \times 10^{-8}$ | 0.000000103 | $2.11 \times 10^{-9}$ | 0.000000101 | $1.93 \times 10^{-9}$ |

**C**
|  |  | Activation<br>Pre vs Post | Inactivation<br>Pre vs Post |
| --- | --- | --- | --- |
| $\mathcal{C}$ | BM | $< 0.001$ | $< 0.001$ |
| | KS | $< 0.001$ | $< 0.001$ |
| $\mathcal{L}$ | BM | $< 0.001$ | $< 0.001$ |
| | KS | $< 0.001$ | $< 0.001$ |
| $\mathcal{V}$ | BM | $< 0.001$ | $< 0.001$ |
| | KS | $< 0.001$ | $< 0.001$ |
| $\eta$ | BM | $< 0.001$ | $< 0.001$ |
| | KS | $< 0.001$ | $< 0.001$ |
| $t_1 - t_0$ | BM | $< 0.001$ | $< 0.001$ |
| | KS | $< 0.001$ | $< 0.001$ |
| $\frac{ds^2}{dt^2}$ | KS | $< 0.001$ | $< 0.001$ |
| $\frac{d}{dt} \frac{ds^2}{dt^2}$ | KS | $< 0.001$ | $< 0.001$ |
| $\left\vert \frac{d}{dt} \frac{ds^2}{dt^2} \right\vert $ | KS | $< 0.001$ | $< 0.001$ |

**Table S4.** The Pearson correlation coefficients and the p-values. (**A**) The Pearson correlation coefficients for the logarithm of information-geometric quantities, the action 𝒞, the length ℒ, the mean velocity 𝒱, the logarithm of the duration *t*_1_ −*t*_0_, and the efficiency *η* in the activation and inactivation processes at the low (L), medium (M), and high (H) cell densities. (**B**) The Pearson correlation coefficients for the logarithm of information-geometric quantities, the action 𝒞, the length ℒ, the mean velocity 𝒱, the logarithm of the duration *t*_1_ −*t*_0_, and the efficiency *η* before (Pre) and after (Post) the Raf inhibitor addition. (**C**) The p-values of no correlation test based on the Pearson correlation coefficients for the conditions, i.e., the low (L), medium (M), and high (H) cell densities, and before (Pre) and after (Post) the Raf inhibitor addition. We calculated the p-values for the logarithm of information-geometric quantities, the action 𝒞, the length ℒ, the mean velocity 𝒱, the logarithm of the duration *t*_1_ −*t*_0_, and the efficiency *η*. The p-values below 0.001 represent as *<* 0.001.

| | $\ln \mathcal{C}$ vs $\ln \mathcal{L}$ | $\ln \mathcal{L}$ vs $\ln \mathcal{V}$ | $\ln \mathcal{V}$ vs $\eta$ | $\ln(t_1 - t_0)$ vs $\ln \mathcal{C}$ | $\ln \mathcal{C}$ vs $\ln \mathcal{V}$ | $\ln \mathcal{L}$ vs $\eta$ | $\ln \mathcal{V}$ vs $\ln(t_1 - t_0)$ | $\eta$ vs $\ln \mathcal{C}$ | $\ln \mathcal{L}$ vs $\ln(t_1 - t_0)$ | $\ln \mathcal{C}$ vs $\ln(t_1 - t_0)$ |
| --- | --- | --- | --- | --- | --- | --- | --- | --- | --- | --- |
| L Activation | 0.925 | 0.612 | -0.0618 | 0.286 | 0.857 | -0.280 | -0.240 | -0.297 | 0.621 | 0.286 |
| L Inactivation | 0.951 | 0.561 | 0.156 | 0.637 | 0.776 | -0.132 | 0.0196 | -0.141 | 0.838 | 0.637 |
| M Activation | 0.934 | 0.679 | -0.154 | 0.195 | 0.889 | -0.403 | -0.269 | -0.395 | 0.525 | 0.195 |
| M Inactivation | 0.944 | 0.616 | 0.0310 | 0.508 | 0.833 | -0.237 | -0.0474 | -0.230 | 0.758 | 0.508 |
| H Activation | 0.921 | 0.622 | -0.0758 | 0.109 | 0.865 | -0.441 | -0.395 | -0.407 | 0.474 | 0.109 |
| H Inactivation | 0.962 | 0.684 | -0.0802 | 0.594 | 0.841 | -0.456 | 0.0772 | -0.445 | 0.780 | 0.594 |

**B**
| | $\ln \mathcal{C}$ vs $\ln \mathcal{L}$ | $\ln \mathcal{L}$ vs $\ln \mathcal{V}$ | $\ln \mathcal{V}$ vs $\eta$ | $\ln(t_1 - t_0)$ vs $\ln \mathcal{C}$ | $\ln \mathcal{C}$ vs $\ln \mathcal{V}$ | $\ln \mathcal{L}$ vs $\eta$ | $\ln \mathcal{V}$ vs $\ln(t_1 - t_0)$ | $\eta$ vs $\ln \mathcal{C}$ | $\ln \mathcal{L}$ vs $\ln(t_1 - t_0)$ | $\ln \mathcal{C}$ vs $\ln(t_1 - t_0)$ |
| --- | --- | --- | --- | --- | --- | --- | --- | --- | --- | --- |
| Pre Activation | 0.978 | 0.894 | -0.334 | 0.315 | 0.966 | -0.402 | 0.0642 | -0.422 | 0.504 | 0.315 |
| Pre Inactivation | 0.987 | 0.907 | -0.0813 | 0.68 | 0.961 | -0.282 | 0.454 | -0.245 | 0.787 | 0.680 |
| Post Activation | 0.973 | 0.851 | -0.441 | 0.570 | 0.945 | -0.455 | 0.277 | -0.516 | 0.740 | 0.570 |
| Post Inactivation | 0.986 | 0.888 | -0.158 | 0.818 | 0.949 | -0.223 | 0.600 | -0.247 | 0.901 | 0.818 |

**C**
| | $\ln \mathcal{C}$ vs $\ln \mathcal{L}$ | $\ln \mathcal{L}$ vs $\ln \mathcal{V}$ | $\ln \mathcal{V}$ vs $\eta$ | $\ln(t_1 - t_0)$ vs $\ln \mathcal{C}$ | $\ln \mathcal{C}$ vs $\ln \mathcal{V}$ | $\ln \mathcal{L}$ vs $\eta$ | $\ln \mathcal{V}$ vs $\ln(t_1 - t_0)$ | $\eta$ vs $\ln \mathcal{C}$ | $\ln \mathcal{L}$ vs $\ln(t_1 - t_0)$ | $\ln \mathcal{C}$ vs $\ln(t_1 - t_0)$ |
| --- | --- | --- | --- | --- | --- | --- | --- | --- | --- | --- |
| L Activation | $< 0.001$ | $< 0.001$ | 0.165 | $< 0.001$ | $< 0.001$ | $< 0.001$ | $< 0.001$ | $< 0.001$ | $< 0.001$ | $< 0.001$ |
| L Inactivation | $< 0.001$ | $< 0.001$ | 0.00155 | $< 0.001$ | $< 0.001$ | 0.00558 | 0.658 | 0.00420 | $< 0.001$ | $< 0.001$ |
| M Activation | $< 0.001$ | $< 0.001$ | $< 0.001$ | $< 0.001$ | $< 0.001$ | $< 0.001$ | $< 0.001$ | $< 0.001$ | $< 0.001$ | $< 0.001$ |
| M Inactivation | $< 0.001$ | $< 0.001$ | 0.0831 | $< 0.001$ | $< 0.001$ | $< 0.001$ | 0.0160 | $< 0.001$ | $< 0.001$ | $< 0.001$ |
| H Activation | $< 0.001$ | $< 0.001$ | $< 0.001$ | $< 0.001$ | $< 0.001$ | $< 0.001$ | $< 0.001$ | $< 0.001$ | $< 0.001$ | $< 0.001$ |
| H Inactivation | $< 0.001$ | $< 0.001$ | $< 0.001$ | $< 0.001$ | $< 0.001$ | $< 0.001$ | $< 0.001$ | $< 0.001$ | $< 0.001$ | $< 0.001$ |
| Pre Activation | $< 0.001$ | $< 0.001$ | $< 0.001$ | $< 0.001$ | $< 0.001$ | $< 0.001$ | 0.0346 | $< 0.001$ | $< 0.001$ | $< 0.001$ |
| Pre Inactivation | $< 0.001$ | $< 0.001$ | 0.0110 | $< 0.001$ | $< 0.001$ | $< 0.001$ | $< 0.001$ | $< 0.001$ | $< 0.001$ | $< 0.001$ |
| Post Activation | $< 0.001$ | $< 0.001$ | $< 0.001$ | $< 0.001$ | $< 0.001$ | $< 0.001$ | $< 0.001$ | $< 0.001$ | $< 0.001$ | $< 0.001$ |
| Post Inactivation | $< 0.001$ | $< 0.001$ | $< 0.001$ | $< 0.001$ | $< 0.001$ | $< 0.001$ | $< 0.001$ | $< 0.001$ | $< 0.001$ | $< 0.001$ |

**Movie S1**.

The fluorescence imaging of the phosphorylated ERK in NRK-52E cells at different cell densities. The movies of the low (left), medium (middle), and high (right) densities are arranged in order from the left. The pseudo-color indicates the FRET/CFP ratio, where the green (red) color shows that the FRET/CFP ratio is 1.0 (2.0) (see also Fig. 2C). The timestamps indicate the time from imaging initiation in minutes.

**Movie S2**.

The fluorescence imaging of the phosphorylated ERK in NRK-52E cells with the Raf inhibitor addition. The Raf inhibitor was added at time 120 min. The pseudo-color indicates the FRET/CFP ratio, where the green (red) color shows that the FRET/CFP ratio is 1.0 (2.0) (see also Fig. 2C). The timestamp indicates the time from imaging initiation in minutes.

